# Differentially acetylated chitosan oligosaccharides as plant defense elicitors against spider mites

**DOI:** 10.1101/2025.10.23.684107

**Authors:** Bowen Jiang, Eryi Ju, Yuguang Du, Bing Wang, Yong Zhao, Jinhua Wei, Takeshi Suzuki, Zhuo A. Wang

## Abstract

The increasing challenges of pesticide resistance and environmental degradation in modern agriculture require sustainable pest management strategies. This study investigates the efficacy of a specific type of hetero-chitooligosaccharide called DACOS, as a plant defense inducer against the two-spotted spider mite, *Tetranychus urticae* Koch. DACOS is characterized with 85% deacetylation and an average molecular weight of 1 kDa. *In planta* bioassays revealed a dose-dependent suppression of mite performance on DACOS-treated leaves of kidney bean seedlings, with peak efficacy occurring at 80 ppm. This concentration reduced mite survival by 30% and fecundity by 59% after six days. In addition to the local effects, mite survival and fecundity were also reduced on untreated leaves of the same seedlings, confirming that the effect of DACOS on plant defense is systemic. Behavioral choice assays revealed that DACOS-treated leaves induced 73% mite repellency. No direct acaricidal effects were observed, which confirms the role of DACOS as a plant defense elicitor. Metabolic profiling revealed enriched pathways (e.g., monoterpenoid biosynthesis and α-linolenic acid metabolism) that produce volatile organic compounds with repellent properties. These findings suggest that DACOS is an environmentally friendly biostimulant that can enhance plant defense, bypass pesticide resistance, and promote the sustainability of crop production.

## Introduction

Modern agriculture is confronted with the dual challenge of increasing productivity to meet global food demand while minimizing its environmental impact on ecosystems and human health. Although synthetic pesticides remain indispensable for protecting crops due to their quick action and high effectiveness against pathogens, pest arthropods and nematodes, weeds, and rodents, decades of overuse of agrochemicals have resulted in soil degradation and loss of biodiversity, necessitating eco-friendly alternatives (Vassilev et al., 2015). These challenges have accelerated the development of sustainable solutions, such as low-toxicity biopesticides and plant biostimulants (Brown and Saa, 2015; Kuc, 2000).

Biostimulants, such as plant defense inducers, have attracted considerable attention. These compounds activate systemic acquired resistance by modulating the immune mechanism without direct pesticidal activity. Notably, marine-derived oligosaccharides have emerged as promising candidates due to their dual functionality in disease resistance and growth regulation, as well as their favorable biodegradability (Yin et al., 2010, 2013). Chitosan oligosaccharide (COS), a marine-derived oligosaccharide, functions as a microbe-associated molecular pattern (MAMP) or a pathogen-associated molecular pattern (PAMP) and importantly, can prime a plant’s immunity like a “vaccine”. By triggering plant defense pathways, it can reduce reliance on conventional pesticides and is a promising biopesticide candidate (Iriti & Faoro, 2009). COS activity depends on its molecular structure, and studies have emphasized that the degree of polymerization, the fraction of acetylation, and even the pattern of acetylation can dramatically alter COS efficacy (Mukarram et al., 2023; Yin et al., 2016). Indeed, many commercial COS mixtures contain both *N*-acetylglucosamine and glucosamine units, and small differences in the acetyl content can switch a response from defense to no effect (Yin et al., 2016). Nonetheless, the literature has broadly documented that COS treatments can enhance disease resistance and sometimes growth across diverse crops by acting as general elicitors of the salicylate, jasmonate, and other defense pathways (El Hadrami et al., 2010).

To address the variability in COS efficacy, we investigated the immune-inducing function of a hetero-chitooligosaccharide (HT-COS) known as DACOS, which has 85% deacetylation and an average molecular weight of 1 kDa. Previous studies have characterized the structural differences of DACOS from two other types of chitooligosaccharides that elicit distinct plant defense: highly acetylated chitin oligosaccharide (composed mainly of *N*-acetylglucosamine) and highly deacetylated COS (composed mainly of glucosamine) (Yin et al., 2016). HT-COSs, which contain *N*-acetylglucosamine and glucosamine in a controlled distribution and polymerization pattern, have strong potential for development as novel biopesticides because they can combine the advantages of both COS and *N*-acetyl-chitooligosaccharide. Specifically, HT-COS have sufficient acetyl groups to bind to plant receptors and effectively activate the plant immune system, but unlike *N*-acetyl-chitooligosaccharide, they maintain good water solubility, even at a high degree of polymerization of more than 8. A structural resolution of DACOS by using nuclear magnetic resonance and mass spectrometry revealed preferential acetylation patterns in long-chain oligomers (Liu et al., 2024). This unique architecture underpins dual bioactivities: one is the aberrant promotion of *Candida tropicalis* proliferation via fungal metabolic reprogramming and biofilm formation (Liu et al., 2024), and the other is enhancing systemic growth in *Brassica napus* through auxin signaling activation and γ-aminobutyric acid accumulation (Tang et al., 2022). However, much remains unknown about the role of DACOS in inducing plant immunity against insect pests.

The two-spotted spider mite, *Tetranychus urticae* Koch (Trombidiformes: Tetranychidae), is a globally distributed phytophagous arthropod, with a broad host range (>1,100 plant species), rapid reproductive capacity, and exceptional environmental adaptability (Migeon et al., 2010; Van Leeuwen, 2015). It severely affects key agricultural crops, such as tomatoes (*Solanum lycopersicum*), strawberries (*Fragaria* × *ananassa*), and cotton (*Gossypium hirsutum*) (Migeon et al., 2010). Current control strategies predominantly rely on synthetic acaricides, including organophosphates and pyrethroids (Zhang et al., 2022), yet they face diminishing returns. Resistance evolution in *T. urticae* has rendered 96 active ingredients ineffective (Dermauw & Van Leeuwen, 2014). Therefore, recent studies have focused on the interaction between mites and plants to improve the defense of plants against *T. urticae* (Santamaria et al., 2020).

The aim of our study is to elucidate the immune effects of DACOS as a plant vaccine in kidney bean (*Phaseolus vulgaris*) against the two-spotted spider mite, focusing on the differences between local and systemic responses and their underlying mechanisms. Here, by using DACOS we demonstrate the induction of resistance and repellency against *T. urticae* in *P. vulgaris*. We evaluated the local (on treated primary leaves) and systemic (on untreated primary leaves paired with treated leaves) effects of DACOS on mite survival, fecundity, feeding activity, and chemo-orientation behavior. In addition, we conducted transcriptome and metabolome analysis to evaluate the effect of DACOS on gene expression and volatile metabolite production for enhancing plant defense mechanisms against *T. urticae*.

## Materials & Methods

### Mites

A *T. urticae* population was collected in London, Ontario, and used for whole-genome sequencing (Grbic et al., 2011). This population was maintained on kidney bean plants *P. vulgaris* at 25 °C under a light period of 16 h/day. Mites were collected using an air-pump system (Cazaux et al., 2014; Suzuki et al., 2017).

### Host plants

Kidney bean seeds ("Hatsumidori No. 2"; Takii, Kyoto, Japan) were placed on a hydrated fir chip substrate (Clean chip; Clea, Tokyo, Japan) and kept under darkness at 25 °C for three days. The germinated seeds were then transplanted into Jiffy-7 pots (Kristiansand, Norway) and cultivated at 25 °C under a 16 h/day light period with a photosynthetic photon flux density of 200 µmol m^−2^ s^−1^. The seedlings were watered every two to three days. After seven days, we sprayed the water solution that has been dissolved with DACOS on the surface of the plant leaves. And seven days after the DACOS application, the leaves were punched using a circular blade with a diameter of 10 mm to obtain leaf discs.

### Sample preparation for gas chromatography–mass spectrometry

The materials were harvested, weighed, immediately frozen in liquid nitrogen, and stored at −80 °C until needed. The samples were ground into a powder in liquid nitrogen. Then, 500 mg of the sample was immediately transferred to a 20 mL headspace vial (Agilent, Palo Alto, CA, USA), containing a 2 mL NaCl saturated solution to inhibit any enzymatic reactions. The vials were then sealed using crimp-top caps with TFE-silicone headspace septa (Agilent). At the time of analysis, each vial was placed in 60 °C for 5 min, then a Smart Solid Phase Microextraction (SPME; Agilent) of 120 µm DVB/CWR/PDMS was exposed to the headspace of the sample for 15 min at 60 °C.

### Quantification of feeding damage

Lesion areas were measured using Adobe Photoshop CC 2020 (Cazaux et al., 2014). Pixel-to-mm calibration was performed using the ruler tool, followed by manual tracing of feeding marks with the pencil tool. Damage was quantified separately for areas treated with water or DACOS solution.

### Mite soaking assay

Adult females three days after final molting were placed on Kimwipes saturated with 0.5% Brilliant Blue FCF solution (control) or 160 ppm DACOS in deionized water. After 1 h, mites were transferred to 10-mm leaf discs (1 mite/disc) from untreated primary leaves of one to two-week-old kidney bean seedlings and kept in polystyrene cups. Leaf discs were replaced three days after transferring the mites. Survival and fecundity were recorded every day over six days.

### Behavioral choice test

Either 80 ppm DACOS or water (control) was applied to the leaves of bean seedlings seven days after germination. Seven days after the foliar application, the leaves were cleaned with water and circular leaf discs (10 mm diameter) were cut out. Cotton was placed in a polypropylene dish and covered with a 90-mm filter paper moistened with water. Leaf discs were then placed on each side of the crossbar of a T-shaped polypropylene sheet and the sheet is above the filter paper. At the beginning of the experiment, one single mite was placed at the bottom end of the T-shaped sheet and allowed to choose between leaf discs on either side. Choice tests were conducted between control and control, control and leaves with local effects (leaves directly treated with DACOS), and control and leaves with systemic effects (untreated leaves paired with treated leaves). The maximum number of trials was set at 28 with different individuals. Mites located outside the leaf disc area after three days were excluded from the data. The number of mite eggs laid on the leaf discs was also recorded.

### Gas chromatography–mass spectrometry

After sampling, desorption of the VOCs from the SPME Arrow coating was carried out in the injection port of the GC apparatus (Model 8890; Agilent) at 250 °C for 5 min. VOCs were identified and quantified in an Agilent Model 8890 GC and a 7000D mass spectrometer (Agilent), equipped with a 30 m × 0.25 mm × 0.25 μm DB-5MS (5% phenyl-polymethylsiloxane) capillary column (Agilent). Helium was used as the carrier gas at a linear velocity of 1.2 mL/min. The injector temperature was kept at 250 °C. The oven temperature was programmed from 40 °C (3.5 min), increasing at 10 °C/min to 100 °C, at 7 °C/min to 180 °C, at 25 °C/min to 280 °C, and kept on hold for 5 min. Mass spectra were recorded in electron impact ionization mode at 70 eV. The quadrupole mass detector, ion source, and transfer line temperatures were set, respectively, at 150, 230, and 280 °C. Mass spectrometry was performed in selected ion monitoring mode to identify and quantify analytes.

### KEGG annotation and enrichment analysis

Identified metabolites were annotated using KEGG Compound database (http://www.kegg.jp/kegg/compound/) and then mapped to KEGG Pathway database (http://www.kegg.jp/kegg/pathway.html).

### Differential metabolites selected

For two-group analysis, differential metabolites were determined by a Variable Importance Projection (VIP) score of VIP > 1 and absolute Log_2_ fold change ≥ 1.0. For multi-group analysis, differential metabolites were determined by VIP (VIP > 1) and *P*-value (P < 0.05, ANOVA). VIP values were extracted from the OPLS-DA result, which also contained score plots and permutation plots, using R package MetaboAnalystR. The data was subjected to log transformation (Log_2_) and mean centering before OPLS-DA. To avoid overfitting, a permutation test (200 permutations) was performed.

### Statistical Analysis

For the fecundity and survival rate, 75 adult mites (three days old) per group were used, 50 mites were used in the soaking test, and 28 mites in the behavioral choice test. These sample sizes provided sufficient detail to meet the study’s objectives. The numbers were chosen to balance robust data collection with ethical considerations, ensuring reliable and meaningful results. Data shown are means ± standard error of the mean. Graphs were generated using GraphPad Prism 7.0. Prior to analysis, data were tested for normality and homogeneity of variance to confirm compliance with the assumptions required for parametric tests. And Statistical significance (*P* < 0.05) was assessed via one-way or two-way ANOVA with Bonferroni post hoc tests for fecundity and leaf damage analysis (SPSS 22.0); survival analysis (Kaplan-Meier) with Log-rank (Mantel-Cox) test (GraphPad Prism 7.0); and statistical analysis for the behavioral choice test using McNemar’s test for number of mites/disc and Mann-Whitney U test for number of eggs/disc (SPSS 22.0). For the KEGG enrichment analysis of differential metabolites, statistical analysis was performed as rich factor = 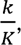 where *k* is the number of differential metabolites that map to the pathway and *K* is the number of those background metabolites that map to the pathway. The *P*-value with hypergeometric test calculated as 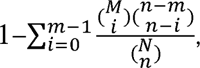 where *N* represents the total number of metabolites with KEGG annotations, *n* represents the number of differential metabolites among *N*, *M* represents the number of metabolites in a certain KEGG pathway among *N*, and *m* represents the number of differentiating metabolites in *M* for a certain KEGG pathway. And statistical analysis for the differential metabolites, the data were standardized by subtracting the mean and dividing by the standard deviation, i.e., 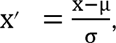 where μ is the mean and σ is the standard deviation. *** *P* < 0.001 and **** *P* <0.0001 indicate statistical differences between the groups. Statistical analysis for volatile metabolites using R studio (MetaboAnalystR) was determined by VIP score and *P*-value (ANOVA).

## Results

### DACOS induced local resistance of kidney bean seedlings to spider mites

Leaves treated with DACOS at concentrations of 16, 80, and 160 ppm reduced mite performance. Six days after transferring the mites onto leaf discs made from treated leaves, the survival declined by 30% and 33% at 80 and 160 ppm, respectively, compared to the control (Fig. 1a). The 16 ppm treatment showed a 7% reduction compared to the control. Cumulative fecundity over six days was inhibited at all concentrations, peaking at 80 ppm with a 59% reduction compared to the control (Fig. 1b). The 16 and 160 ppm treatments reduced fecundity by 46% and 33%, respectively, compared to the control. Significant inhibition of fecundity was observed only at 80 ppm one day after transferring the mites (Fig. 1c).

**Fig. 1.**
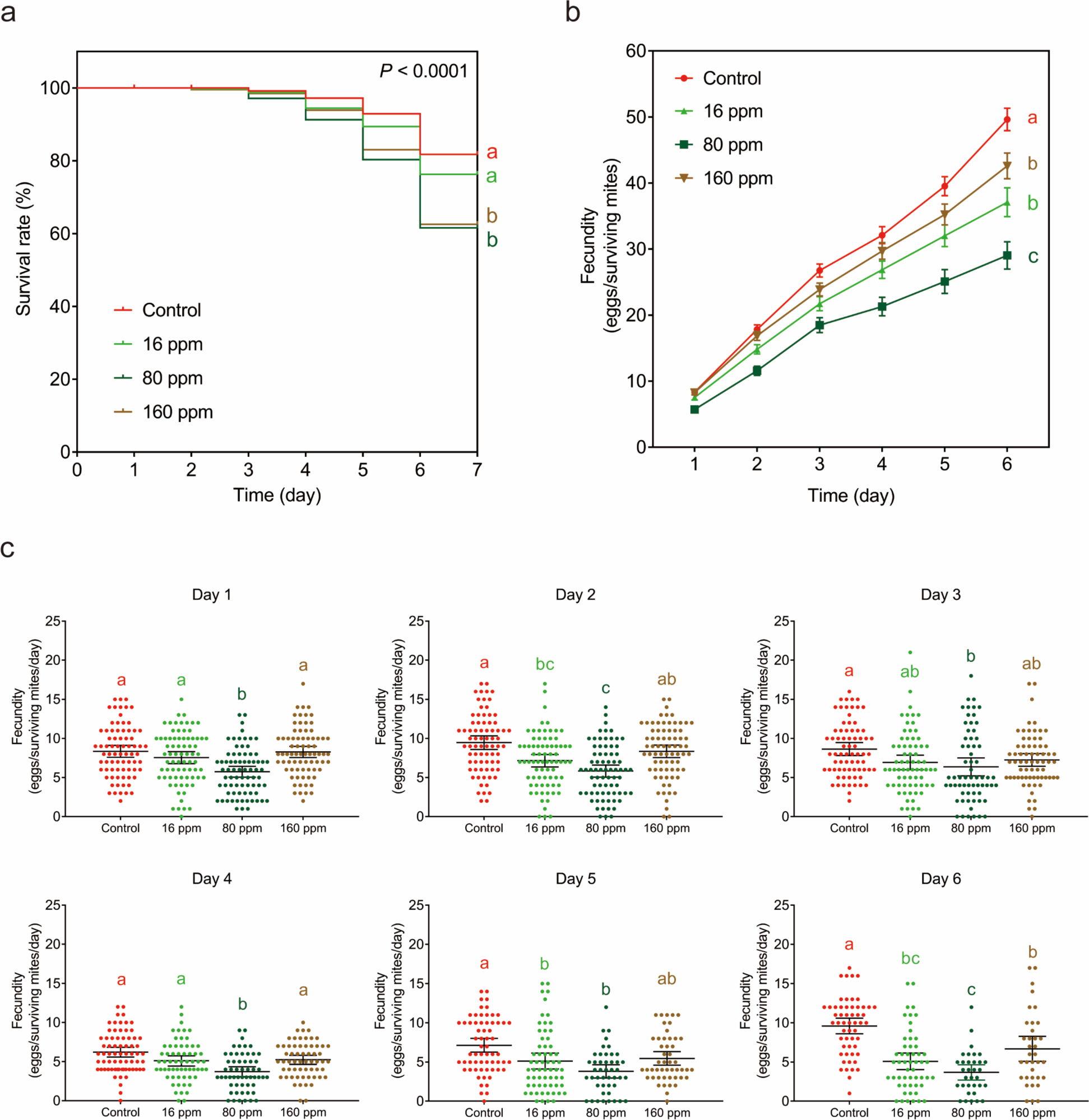
Plant-mediated local effects of DACOS treatment on the performance of spider mites. Adult females of *Tetranychus urticae* were introduced onto kidney bean leaf discs made from primary leaves treated with 16, 80, or 160 ppm DACOS or 0 ppm (control). Mite performance was evaluated by (a) survival, (b) cumulative fecundity, and (c) daily fecundity. Survival curves were plotted using the Kaplan-Meier method, and statistical differences were analyzed using the log-rank test (*P* < 0.0001). Data on the cumulative and daily fecundity are presented as mean ± SEM. Additionally, individual data were plotted for the daily fecundity. Cumulative and daily fecundity were statistically analyzed using a one-way ANOVA followed by a Bonferroni’s multiple comparison test. Data with different letters were statistically significant at *P* < 0.05

### DACOS induced systemic resistance of kidney bean seedlings to spider mites

To elucidate systemic resistance, a bioassay was conducted using paired untreated leaves adjacent to the primary leaves treated with DACOS on the same seedlings. The systemic effect of DACOS reduced mite performance at all concentrations, peaking at 160 ppm. Six days after transferring the mites onto the leaf discs, the survival declined by 19%, 11%, and 28% at 16, 80, and 160 ppm, respectively, compared to the control (Fig. 2a). Cumulative fecundity over six days was inhibited by 42%, 41%, and 33% at 16, 80, and 160 ppm, respectively, compared to the control (Fig. 2b). Significant inhibition of fecundity was observed only at 160 ppm one day after transferring the mites (Fig. 2c). Six days after transferring the mites, fecundity decreased by 42%, 39%, and 62% in the DACOS treatments at 16, 80, and 160 ppm, respectively, compared to the control.

**Fig. 2.**
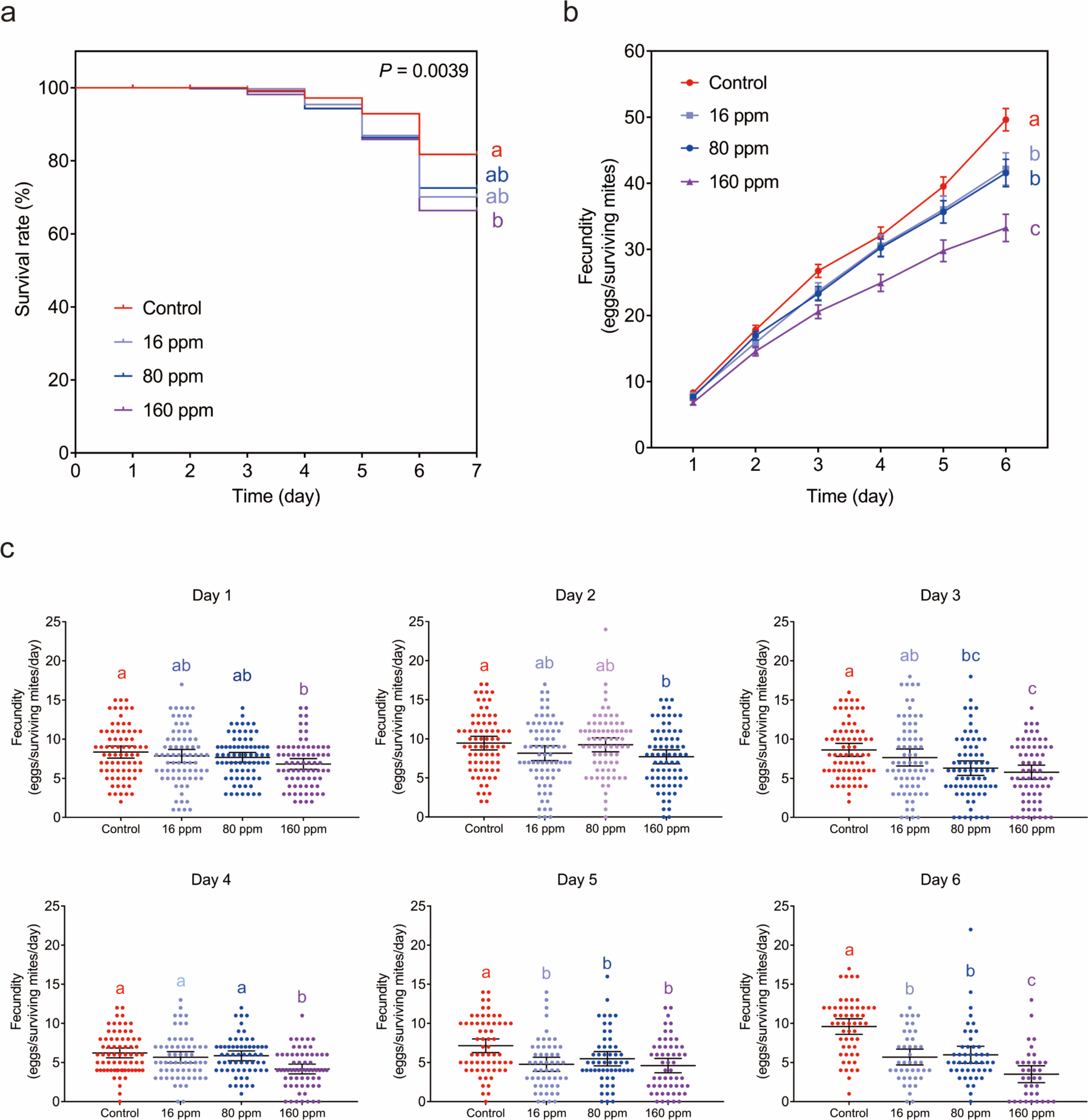
Plant-mediated systemic effects of DACOS treatment on the performance of spider mites. Adult females of *Tetranychus urticae* were introduced onto kidney bean leaf discs from untreated primary leaves of seedlings whose other primary leaves were treated with 16, 80, or 160 ppm DACOS or 0 ppm (control). Mite performance was evaluated by (a) survival, (b) cumulative fecundity, and (c) daily fecundity. Survival curves were plotted using the Kaplan-Meier method, and statistical differences were analyzed using the log-rank test (*P* < 0.0001). Data on the cumulative and daily fecundity are presented as mean ± SEM. Additionally, individual data were plotted for the daily fecundity. Cumulative and daily fecundity were statistically analyzed using a one-way ANOVA followed by a Bonferroni’s multiple comparison test. Data with different letters are statistically significant at *P* < 0.05

### DACOS induced local and systemic effects on mite feeding in kidney bean seedlings

The local and systemic effects of the DACOS treatment significantly reduced the feeding damage caused by spider mites (Fig. 3a). The local effect of the 80-ppm DACOS treatment reduced the area of chlorotic spots by 58% compared to the control, with the 16- and 160-ppm treatments reducing the area by 31% and 36%, respectively (Fig. 3b). The systemic effect of the 160-ppm DACOS treatment reduced the area of chlorotic spots by 54% compared to the control with the 16- and 80-ppm treatments reducing the area by 30% and 41%, respectively (Fig. 3c). Notably, a physiological disorder (see Fig. 3a) occurred in leaf discs made from leaves treated with 160 ppm DACOS, likely reflecting immune over-activation.

**Fig. 3.**
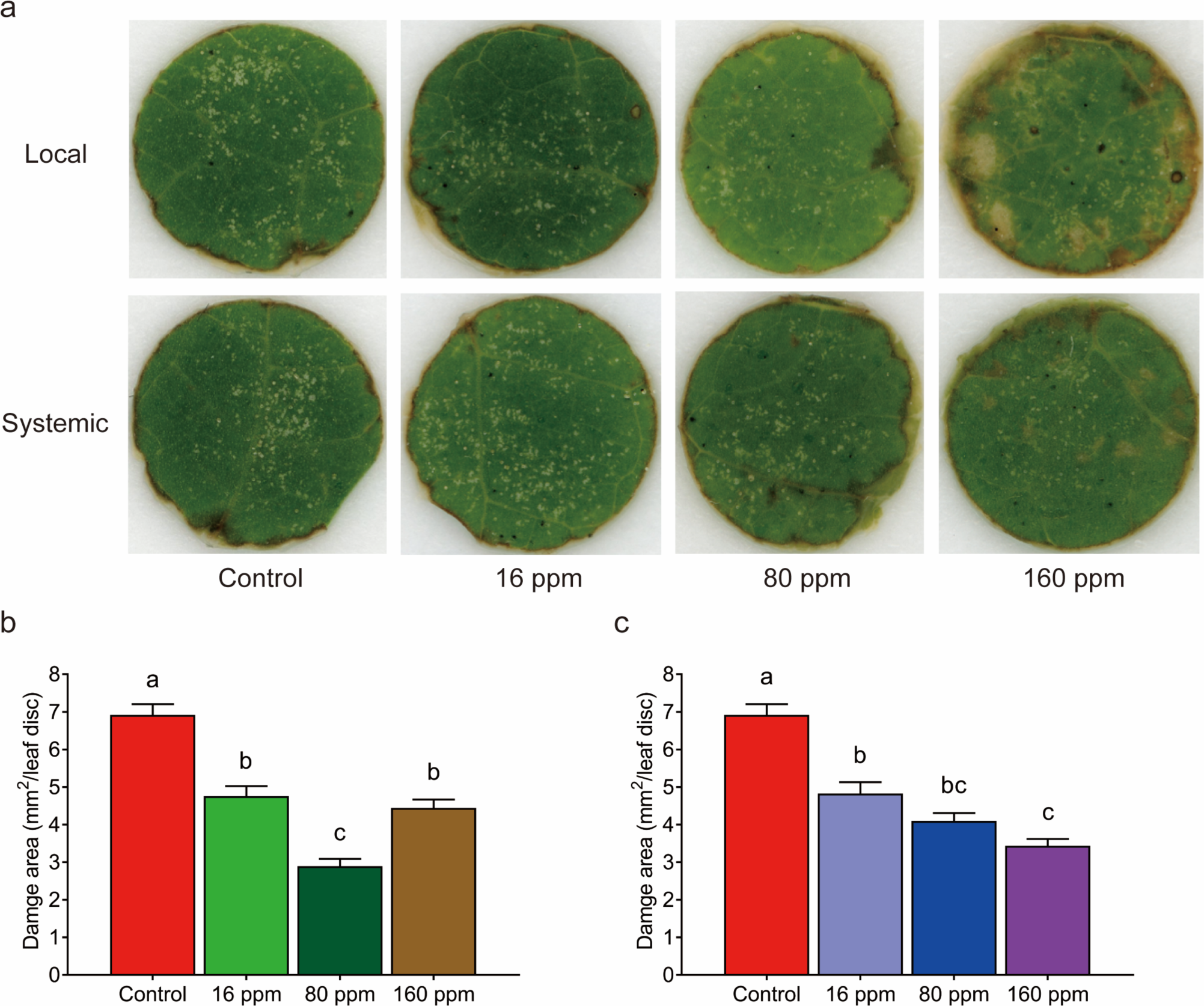
Comparison of plant-mediated local and systemic effects of DACOS treatment on mite feeding. (a) Chlorotic spots caused by mite feeding on leaf discs that were treated with DACOS directly (local) or indirectly (systemic). (b, c) Quantification of chlorotic spots on leaf discs from locally (b) and systemically (c) treated leaves. Data are presented as mean ± SEM. Feeding area was statistically analyzed using a one-way ANOVA followed by a Bonferroni’s multiple comparison test. Data with different letters are statistically significant at *P* < 0.05

### Direct effects of DACOS on spider mite performance

No significant differences in survival (Fig. 4a) and fecundity (Fig. 4b) were observed in mites soaked in 160 ppm DACOS for 1 h compared to the control. However, the feeding activity of mites soaked in 160 ppm DACOS was 30% greater than that of the control (Figs. 4c and 4d).

**Fig. 4.**
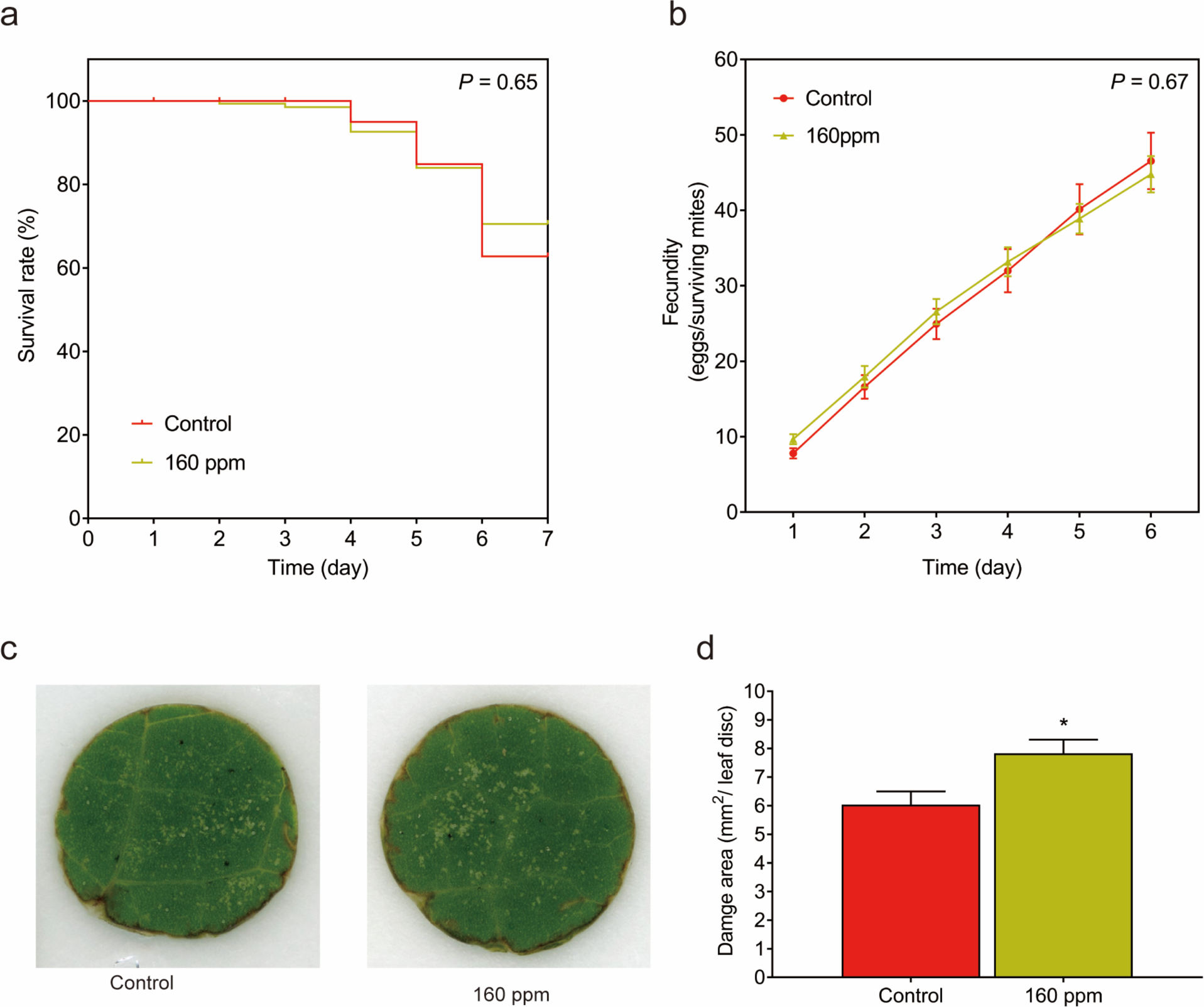
Direct effects of DACOS on the performance of spider mites. (a) Survival curves were plotted using the Kaplan-Meier method, and statistical differences were analyzed using the log-rank test (*P* > 0.05). (b) Data on the cumulative fecundity are presented as mean ± SEM. (c) Chlorotic spots caused by mite feeding on leaf discs. (d) Quantification of the chlorotic spots on leaf discs. Cumulative fecundity and feeding area were statistically analyzed using a one-way ANOVA followed by a Bonferroni’s multiple comparison test. Data with different letters are statistically significant at *P* < 0.05

### Chemo-orientation behavior of spider mites toward DACOS-treated leaves

Behavioral choice assays revealed mite repellency in DACOS-treated leaves. The 80-ppm treatment reduced mite localization by 73% and 58% one and two days, respectively, after starting the assay (Fig. 5a). Three days after starting the assay, differences in mite localization diminished. Fecundity per female decreased by 54%, 52%, and 54% after one, two, and three days, respectively, of starting the assay (Fig. 5b).

**Fig. 5.**
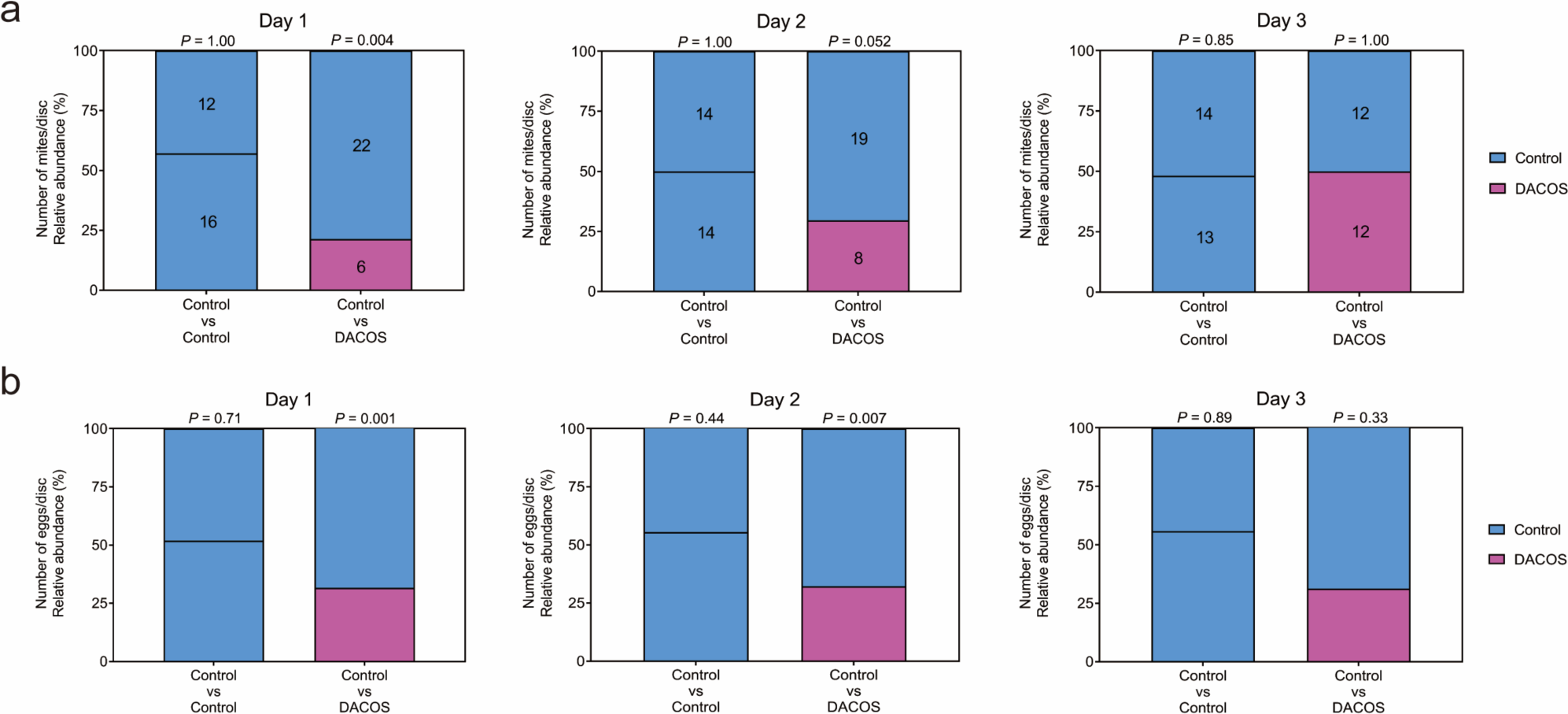
Behavioral choice test for evaluating mite selection of DACOS-treated leaf discs. In a T-shaped choice test, each adult female mite was allowed over a period of 3 days to select an untreated leaf disc (control) or a leaf disc treated with 80 ppm DACOS. The proportions of (a) the number of surviving mites and (b) the number of eggs laid per surviving mite on each leaf disc were plotted (*n* = 28). Dead mites and mites who were not located on leaf discs were omitted from the analysis. Statistical analysis was performed using (a) McNemar’s test and (b) Mann-Whitney U test (for 2 groups)

### Local and systemic gene expression responses of kidney bean seedlings to DACOS

Since the suppressive effects of DACOS application on mite performance were greater locally than systemically, we further dissected their transcriptomic differences. On average across three biological replicates, bean leaves produced approximately 7.36 Gb of transcriptome reads for local responses and 8.07 Gb for systemic responses (Table S1). Untreated (control) bean leaves produced approximately 8.25 Gb.

A principal component analysis of the gene expression profiles revealed distinct clustering of local, systemic, and control samples (Fig. 6a). Differential expression analysis identified numerous differentially expressed genes (DEGs), as visualized in Figure 6b. As shown in Figure S1a, in local and systemic samples a total of 1,460 and 1,570 genes were found to be up- and down-regulated, respectively, compared to the control, of which the DEGs in local samples compared to control were 3,373 (1,607 up-regulated and 1,766 down-regulated) and DEGs in systemic samples were 2,685 (1,214 up-regulated and 1,471 down-regulated). The comparison between samples responding systemically and those responding locally to DACOS identified 490 DEGs (27 up-regulated and 463 down-regulated). The KEGG pathway enrichment analysis of the DEGs in local and systemic samples compared with the control identified the top 50 enriched pathways, which were predominantly in the metabolism and organismal systems categories and included plant hormone signal transduction, MAPK signaling, phenylpropanoid biosynthesis, and flavonoid biosynthesis (Fig. 6c). In local samples compared to systemic samples, KEGG enrichment of DEGs occurred mainly in metabolism and environmental information processing (Fig. S1b). Enriched pathways included the MAPK signaling pathway, plant hormone signal transduction, flavonoid biosynthesis, and phenylpropanoid biosynthesis. In addition, the DEGs enriched in Gene Ontology included molecular function, biological processes, and cellular components (Fig. 6d), which suggests that resources were reallocated from growth to defense. However, Gene Ontology enrichment showed a widespread downregulation in systemic versus local comparisons (Fig. S1c).

**Fig. 6.**
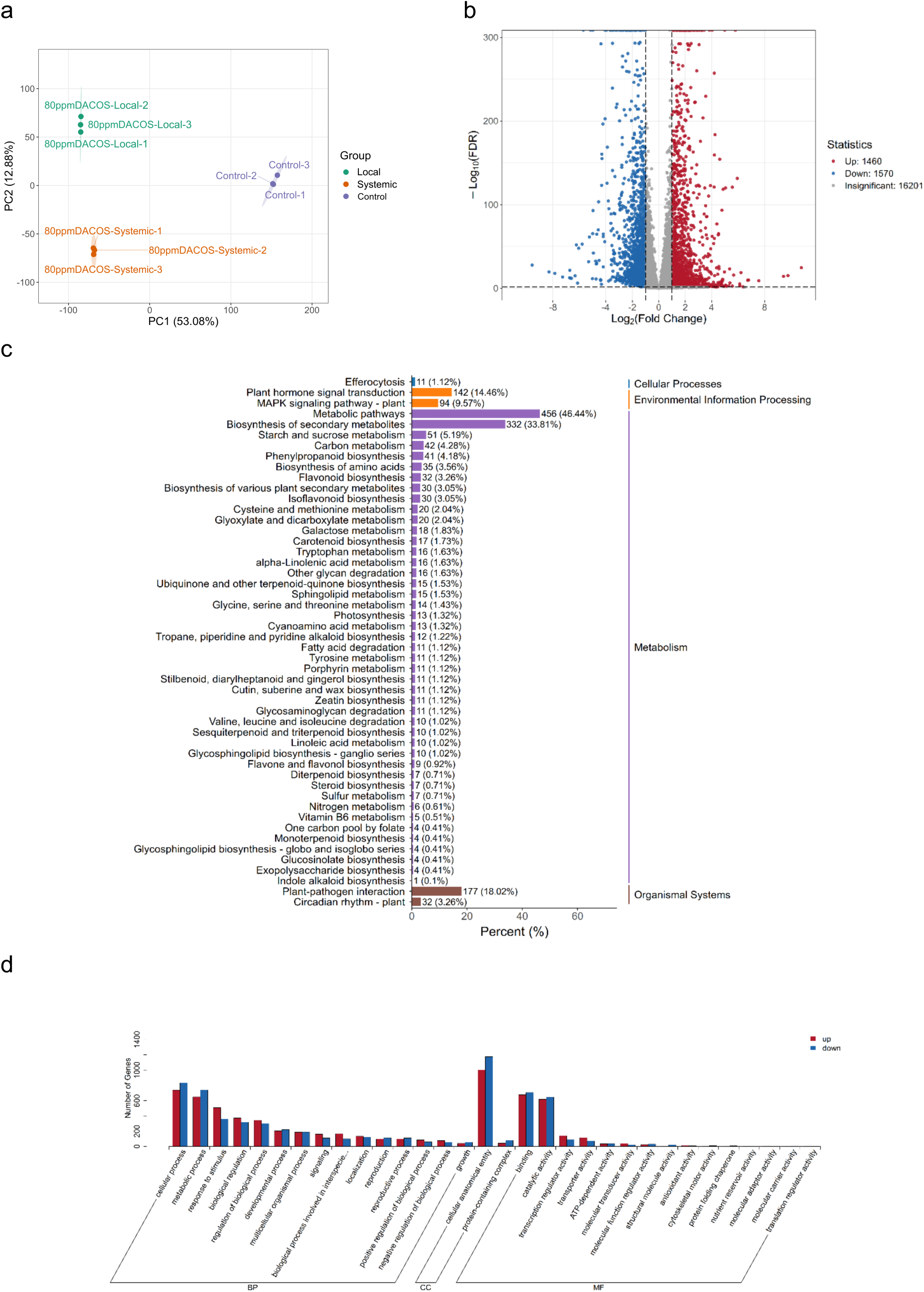
Gene expression of *Phaseolus vulgaris* between control (water) and DACOS treatments. (a) The principal component (PC) analysis in gene expression. (b) A volcano plot of differentially expressed genes. Red plots indicate DACOS is significantly higher than that of the control, and blue plots indicate the opposite. (c) A KEGG pathway enrichment of differentially expressed genes. The x-axis represents the number of differentially expressed genes annotated to the pathway, and the y-axis represents the name of the KEGG pathway. The actual number of differentially expressed genes is noted beside each pathway, and the values in parentheses are the ratios to the total number of differentially expressed genes. The labels on the right give the KEGG pathway classification. (d) The Genes Ontology (GO) enrichment analysis of significantly up-regulated genes. The x-axis represents the secondary GO entries categorized into biological processes (BP), cellular components (CC), and molecular function (MF), and the y-axis represents the number of differentially expressed genes for each GO entry

### DACOS induces the production of plant defense–related metabolites

To complement transcriptomic analyses and uncover the biochemical mechanisms driving DACOS-elicited resistance to spider mites, gas chromatography–mass spectrometry was employed to profile volatile metabolites in kidney bean leaves. Comparisons encompassed DACOS treatment versus control (water), as well as local versus systemic effects, given the superior resistance efficacy of local application.

Orthogonal partial least squares discriminant analysis (OPLS-DA) of metabolite profiles demonstrated a clear separation of DACOS-treated samples from the control (Fig. S2a). KEGG enrichment of differential metabolites in the DACOS-treated samples versus the control highlighted the defense pathways, including arginine biosynthesis, purine metabolism, and α-linolenic acid metabolism (Fig. 7a), and difference in the systemically vs locally treated samples, including monoterpenoid biosynthesis, and α-linolenic acid metabolism (Fig. S2b). Volcano plots comparing the effects of DACOS treatments showed more up-regulated metabolites in local versus control than in systemic versus control, and more down-regulated metabolites in systemic versus local (Fig. S2c). Heatmap clustering segregated the treatments, with local and systemic responses to DACOS treatment elevating D-carvone and (-)-carvone, while control samples showed depletions of, for example, higher D-limonene, cyclohexene and L-menthone (Fig. 7b). Z-score plots confirmed positive shifts in local treatment with DACOS compared to systemic and control for jasmonates and terpenoids (Fig. 7c and Table S2). The significant contributions of general metabolic pathways and secondary metabolite biosynthesis (66% and 38%, respectively) highlight the extensive metabolic reprogramming in local compared to systemic responses to DACOS treatment (Fig. S2d). Over 20 differentially abundant volatile metabolites were detected, including four terpene species, four hydrocarbons, three esters, two phenolic compounds, and one each of aldehydes, ketones, alcohols, acids, and sulfur-containing compounds (Figs. 7d and S3). Monoterpenes can act as direct toxins or repellents to insects and attract natural enemies, such as parasitoids and predators, thereby providing indirect defense (Fontana et al., 2009). To integrate transcriptomic and metabolomic datasets, DEGs and differential volatile metabolites were mapped onto the KEGG pathway ko00592 for α-linolenic acid metabolism (Fig. S4). A parallel branch to green leaf volatiles showed activation via hydroperoxide lyase, leading to aldehydes such as 3-hexenal and alcohols like 3-hexen-1-ol, with a differential metabolite highlighted at 3-hexen-1-ol acetate. (Fig. S4).

**Fig. 7.**
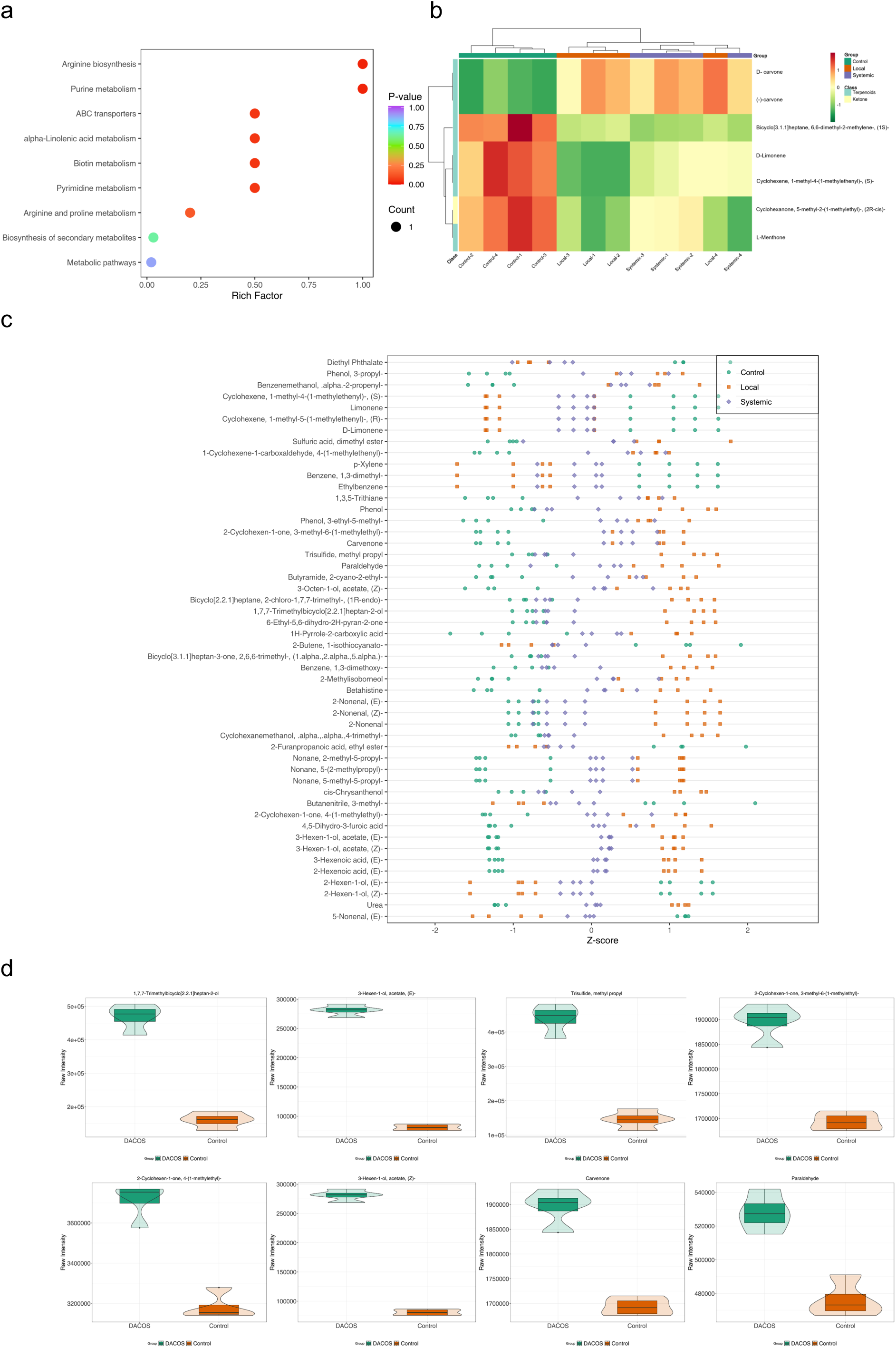
Volatile metabolites induced in leaves treated with DACOS. (a) KEGG enrichment analysis of differential metabolites. The color of the points indicates the magnitude of the *P*-value, with red representing a more significant enrichment. Each pathway was enriched by one differentially expressed metabolite. (b) A heat map of metabolite changes. Different colors represent the different values obtained after standardization of relative content (red: high, green: low). Color-coding above the heatmap corresponds to the treatment concentration (Group), the tree diagram on the left of the heatmap represents the hierarchical clustering results of the differential metabolites, with color-coding used for the first-level classification of substances (Class). (c) Z-score plot of differential metabolites. Types of treatment are shown in different colors for each of the top 50 metabolites ranked by variable importance. (d) Quantification of the differential metabolites. The raw intensity indicates the peak signal area of each metabolite. Darker-colored boxes indicate the interquartile range, whisker lines the 95% confidence interval, and horizontal lines the median. Lighter-colored shapes represent the distribution density of the data

### DACOS effects on the growth of kidney bean seedlings

Foliar application of DACOS demonstrated no effect on shoot growth (Figs. 8a–c, e); however, it promoted the root growth (Figs. 8d, S5). Six days after foliar application of DACOS at 16, 80, and 160 ppm, root dry weights were 1.5, 1.9, and 1.7 times higher than those in the control, respectively.

**Fig. 8.**
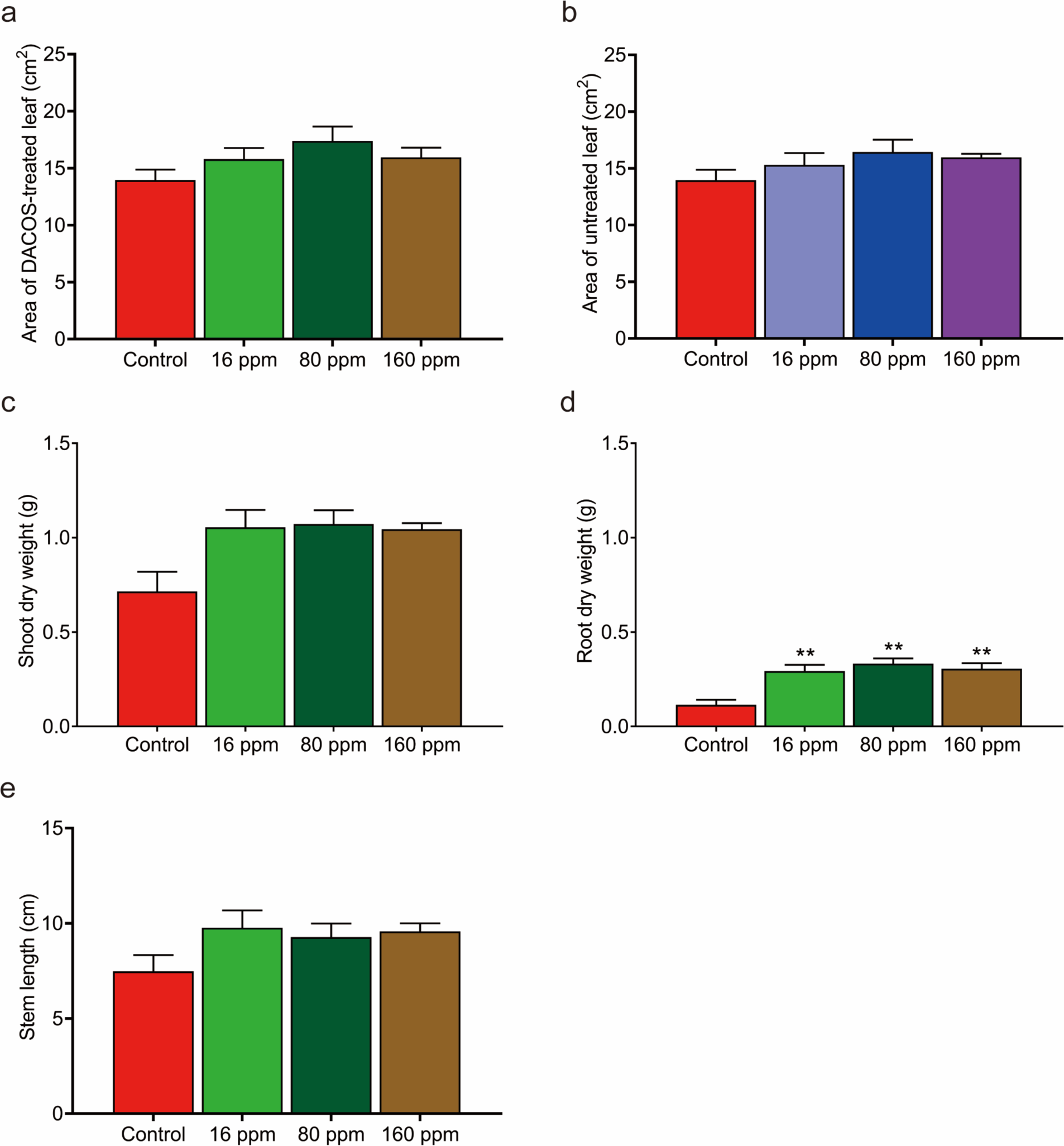
Effect of DACOS treatment on plant growth. Kidney bean plants were treated with 16, 80, or 160 ppm DACOS or 0 ppm (control). Plant growth was evaluated by measuring the leaf area of (a) locally and (b) systemically treated plants, the dry weight of (c) shoots and (d) roots, and (e) the stem length. Data on plant growth are presented as the mean ± SEM. Plant growth was statistically analyzed using a Student’s independent *t* test and found no significance between groups

## Discussion

The study revealed that DACOS functions as a plant immune inducer, triggering local and systemic resistance in kidney bean seedlings and significantly reducing the survival, fecundity, and feeding activity of spider mites. Foliar application of DACOS induced local and systemic responses in bean seedlings, inhibiting the survival, fecundity, and feeding activity of spider mites (Figs. 1, 2, and 3). Local and systemic responses reached their peak at concentrations of 80 and 160 ppm of DACOS, respectively. This aligns with the "optimal dosage effect" observed in plant immune responses, whereby excessive levels of elicitors can lead to cytotoxicity or immunosuppression (van Hulten et al., 2006). The inhibitory effects on mite fecundity persisted for at least six days (Figs. 1 and 2). Behavioral choice assays and omics analysis revealed that plant-derived volatile metabolites derived from bean leaves treated with DACOS may contribute to mite repellency (Figs. 5, 6, and 7). Furthermore, stronger enrichment occurred in local effects (Fig. S2b), emphasizing terpenoid volatile organic compounds (VOCs), including volatile monoterpenoids, jasmonates, and alkaloids for mite repellency. Consequently, DACOS elicits induced resistance by upregulating defense-related pathways, genes, and metabolites. Local effects are more effective than systemic effects, thereby necessitating higher doses for optimal systemic responses to achieve comparable mite repellence (Figs. 6 and 7).

Previous studies have demonstrated that COS, acting as MAMPs, trigger plant innate immunity through both direct antifungal activity (by suppressing spore germination) and induced systemic resistance mechanisms (Narula et al., 2020). Notably, COS reduced the infectivity of the cucumber mosaic virus by 45% in passionfruit (*Passiflora edulis*) (L. Zhang et al., 2023). In grapes (*Vitis vinifera*), chitosan with a low degree of polymerization (DP = 12–20) in conjunction with fungicides achieved 47% to 54% efficacy in controlling powdery mildew (*Erysiphe necator*), comparable to the 73% to 80% efficacy of conventional copper-based treatments (Brule et al., 2024). Our findings expand the COS paradigm to include plant resistance against herbivorous arthropods. From an agricultural standpoint, the increasing resistance of spider mites to pesticides poses a significant threat to global crop production (Ghazy et al., 2019). DACOS mitigates the evolution of resistance by leveraging plant-mediated "indirect control" via immune activation. DACOS induces repellent volatiles, such as monoterpenes and jasmonic acid, that disrupt host-seeking behavior in spider mites (Figs. 6 and 7) while attracting generalist predators (Mukhtar Ahmed et al., 2020; War et al., 2012). And particularly, Jasmonic acid is derived from α-linolenic acid through a series of enzymatic reactions in the chloroplast and peroxisome. This leads to the activation of defense-related genes and the production of secondary metabolites (Wasternack & Hause, 2013). Jasmonic acid signaling is particularly important for responses to chewing herbivorous arthropods because it triggers the synthesis of proteinase inhibitors and other defensive compounds (Wasternack & Hause, 2013). Although this tritrophic interaction still requires empirical validation, the absence of direct acaricidal activity (Figs. 4a and 4b) minimizes non-target risks and supports ecological sustainability.

DACOS induced local anti-mite responses with optimal efficacy at 80 ppm (Fig. 1). Mites damage crops by piercing mesophyll cells and extracting cellular contents, which leads to chloroplast disintegration and photosynthetic impairment (Bensoussan et al., 2016). COS modulate plant immune responses by activating the brassinosteroid signaling pathway and ROS/NO/eATP signaling cascades in a coordinated manner (Narula et al., 2020). The volatile metabolites identified in our study, particularly borneol and 3-hexen-1-ol acetate, are well documented for their roles in plant defense (Scala et al., 2013). Borneol’s repellent properties make it a key player in direct defense, potentially reducing insect feeding and fecundity on kidney bean seedlings. Similarly, green leaf volatiles such as 3-hexen-1-ol acetate contribute to indirect defense by attracting natural enemies that can reduce herbivore populations. The sulfur compound methyl propyl trisulfide enhances direct defense through its repellent odor.

A previous study indicated that chitosan effectively altered secondary metabolites, increasing alkaloids by 38% and terpenoids by 23% (Ziaaddini et al., 2022). Here, we conducted a behavioral choice assay to show that DACOS treatment may prompt plants to release volatile metabolites for pest defense (Fig. 7). The temporal dynamics of the repellency induced by DACOS treatment warrant particular attention. At a concentration of 80 ppm, DACOS caused 73% and 58% reductions in mite colonization on Days 1 and 2, respectively (Figs. 5a and 5b). This suggests that DACOS induces the emission of VOCs, particularly monoterpenoids and green leaf volatiles, which decrease mite performance (Fontana et al., 2009; Scala et al., 2013). However, diminished repellency by Day 3 (Fig. 5a) coincided with the depletion of metabolites observed in chitosan-treated grapevines, where the peak VOC emission was detected at 48 h (Brule et al., 2024). This highlights the dynamic temporal regulation of these metabolites and necessitates further investigation into their synthesis and degradation pathways. Additionally, the divergence in localized and systemic effects at different concentrations (optimal concentration of 160 ppm for systemic tissues but 80 ppm for local tissues; Figs. 1 and 2) suggests that there are distinct signaling thresholds for the propagation of immune responses within and between tissues.

Beyond pest resistance, COS promotes growth in *Brassica napus* via modulation of the Aux/IAA gene (Tang et al., 2022) and improves drought tolerance in maize by balancing superoxide dismutase/peroxidase activities (Li et al., 2022). Our results showed DACOS treatment had no side effect on plant growth, and significantly promoted root growth. These synergies suggest that COS strengthens plant immune barriers and optimizes photoassimilate allocation (via root-shoot signaling) and antioxidant systems, providing dual "resilience–growth" benefits (Li et al., 2022).

In conclusion, DACOS is a novel plant immune inducer that may facilitate innovative solutions for the sustainable control of spider mites by eliciting systemic acquired resistance and inducing the production of VOCs in plants. This approach circumvents pesticide resistance while preserving ecological balance. Future studies should use metabolomics to profile the secondary metabolites induced by DACOS (e.g., terpenoids and alkaloids) and clarify how they modulate the behavior of spider mites. Field trials integrating DACOS with biocontrol agents could validate its compatibility with Integrated Pest Management guidelines. Additionally, nanoformulations could extend its functional longevity (Saberi Riseh et al., 2024). Due to its concentration-dependent efficacy, environmental safety, and dual-action synergy with plant growth pathways, DACOS is a promising candidate for next-generation sustainable agriculture.

## Statements & Declarations

### Data availability

The datasets generated during and/or analyzed during the current study are available from the corresponding author on reasonable request.

### Acknowledgement

We thank Faten Abdelsalam Hamdi for the T-Shaped method. We thank Hiroki Kitara for the fecundity and soaking test methods. We thank Hideaki Takahashi for the method of detecting volatile metabolites.

### Author contributions

BWJ: data collection, statistical analyses, data interpretation, and the initial draft of the manuscript; EJ: analyses and interpretation of the results; YD, BW, YZ, JW: comments and revisions on the manuscript; TS: conceptualization and design of the study, analyses and interpretation of the results, final writing; ZAW, conceptualization and design of the study.

### Competing interests

None of the authors have any conflict of interest related to the study.

### Funding

This work was supported partly by the JSPS KAKENHI (24K21256) to TS and the National Key R&D Program of China (2024YFD2402105) to ZAW.

## Notes

### Competing Interest Statement

The authors have declared no competing interest.

## References

Bensoussan, N., Santamaria, M. E., Zhurov, V., Diaz, I., Grbic, M., & Grbic, V. (2016). Plant-Herbivore Interaction: Dissection of the Cellular Pattern of Tetranychus urticae Feeding on the Host Plant. Front Plant Sci, 7, 1105. doi:10.3389/fpls.2016.01105

Brown, P., & Saa, S. (2015). Biostimulants in agriculture. Front Plant Sci, 6, 671. doi:10.3389/fpls.2015.00671

Brule, D., Heloir, M. C., Roudaire, T., Villette, J., Bonnet, S., Pascal, Y., . . . Poinssot, B. (2024). Increasing vineyard sustainability: innovating a targeted chitosan-derived biocontrol solution to induce grapevine resistance against downy and powdery mildews. Front Plant Sci, 15, 1360254. doi:10.3389/fpls.2024.1360254

Cazaux, M., Navarro, M., Bruinsma, K. A., Zhurov, V., Negrave, T., Van Leeuwen, T., . . . Grbic, M. (2014). Application of two-spotted spider mite Tetranychus urticae for plant-pest interaction studies. J Vis Exp(89). doi:10.3791/51738

Dermauw, W., & Van Leeuwen, T. (2014). The ABC gene family in arthropods: comparative genomics and role in insecticide transport and resistance. Insect Biochem Mol Biol, 45, 89–110. doi:10.1016/j.ibmb.2013.11.001

El Hadrami, A., Adam, L. R., El Hadrami, I., & Daayf, F. (2010). Chitosan in plant protection. Mar Drugs, 8(4), 968–987. doi:10.3390/md8040968

Fontana, A., Reichelt, M., Hempel, S., Gershenzon, J., & Unsicker, S. B. (2009). The effects of arbuscular mycorrhizal fungi on direct and indirect defense metabolites of Plantago lanceolata L. J Chem Ecol, 35(7), 833–843. doi:10.1007/s10886-009-9654-0

Ghazy, N. A., Gotoh, T., & Suzuki, T. (2019). Impact of global warming scenarios on life-history traits of Tetranychus evansi (Acari: Tetranychidae). BMC Ecol, 19(1), 48. doi:10.1186/s12898-019-0264-6

Grbic, M., Van Leeuwen, T., Clark, R. M., Rombauts, S., Rouze, P., Grbic, V., . . . Van de Peer, Y. (2011). The genome of Tetranychus urticae reveals herbivorous pest adaptations. Nature, 479(7374), 487–492. doi:10.1038/nature10640

Iriti, M., & Faoro, F. (2009). Chitosan as a MAMP, searching for a PRR. Plant Signal Behav, 4(1), 66–68. doi:10.4161/psb.4.1.7408

Kuc, J. (2000). Development and future direction of induced systemic resistance in plants. Crop Protection, 19(8-10), 859–861. doi:Doi 10.1016/S0261-2194(00)00122-8

Li, J., Han, A., Zhang, L., Meng, Y., Xu, L., Ma, F., & Liu, R. (2022). Chitosan oligosaccharide alleviates the growth inhibition caused by physcion and synergistically enhances resilience in maize seedlings. Sci Rep, 12(1), 162. doi:10.1038/s41598-021-04153-3

Liu, Y., Li, R., Zhang, Y., Jiao, S., Xu, T., Zhou, Y., . . . Wang, Z. A. (2024). Unveiling the inverse antimicrobial impact of a hetero-chitooligosaccharide on Candida tropicalis growth and biofilm formation. Carbohydr Polym, 333, 121999. doi:10.1016/j.carbpol.2024.121999

Migeon, A., Nouguier, E., & Dorkeld, F. (2010). Spider Mites Web: A comprehensive database for the Tetranychidae. Trends in Acarology, 557-560. doi:10.1007/978-90-481-9837-5_96

Mukarram, M., Ali, J., Dadkhah-Aghdash, H., Kurjak, D., Kacik, F., & Durkovic, J. (2023). Chitosan-induced biotic stress tolerance and crosstalk with phytohormones, antioxidants, and other signalling molecules. Front Plant Sci, 14, 1217822. doi:10.3389/fpls.2023.1217822

Mukhtar Ahmed, K. B., Khan, M. M. A., Siddiqui, H., & Jahan, A. (2020). Chitosan and its oligosaccharides, a promising option for sustainable crop production- a review. Carbohydr Polym, 227, 115331. doi:10.1016/j.carbpol.2019.115331

Narula, K., Elagamey, E., Abdellatef, M. A. E., Sinha, A., Ghosh, S., Chakraborty, N., & Chakraborty, S. (2020). Chitosan-triggered immunity to Fusarium in chickpea is associated with changes in the plant extracellular matrix architecture, stomatal closure and remodeling of the plant metabolome and proteome. Plant J, 103(2), 561–583. doi:10.1111/tpj.14750

Saberi Riseh, R., Vatankhah, M., Hassanisaadi, M., & Varma, R. S. (2024). A review of chitosan nanoparticles: Nature’s gift for transforming agriculture through smart and effective delivery mechanisms. Int J Biol Macromol, 260(Pt 2), 129522. doi:10.1016/j.ijbiomac.2024.129522

Santamaria, M. E., Arnaiz, A., Rosa-Diaz, I., Gonzalez-Melendi, P., Romero-Hernandez, G., Ojeda-Martinez, D. A., . . . Diaz, I. (2020). Plant Defenses Against Tetranychus urticae: Mind the Gaps. Plants (Basel), 9(4). doi:10.3390/plants9040464

Scala, A., Allmann, S., Mirabella, R., Haring, M. A., & Schuurink, R. C. (2013). Green leaf volatiles: a plant’s multifunctional weapon against herbivores and pathogens. Int J Mol Sci, 14(9), 17781–17811. doi:10.3390/ijms140917781

Suzuki, T., Espana, M. U., Nunes, M. A., Zhurov, V., Dermauw, W., Osakabe, M., . . . Grbic, V. (2017). Protocols for the delivery of small molecules to the two-spotted spider mite, Tetranychus urticae. PLoS One, 12(7), e0180658. doi:10.1371/journal.pone.0180658

Tang, C., Zhai, Y., Wang, Z., Zhao, X., Yang, C., Zhao, Y., . . . Zhang, D. Y. (2022). Metabolomics and transcriptomics reveal the effect of hetero-chitooligosaccharides in promoting growth of Brassica napus. Sci Rep, 12(1), 21197. doi:10.1038/s41598-022-25850-7

van Hulten, M., Pelser, M., van Loon, L. C., Pieterse, C. M., & Ton, J. (2006). Costs and benefits of priming for defense in Arabidopsis. Proc Natl Acad Sci U S A, 103(14), 5602–5607. doi:10.1073/pnas.0510213103

Van Leeuwen, T., Tirry, L., Yamamoto, A., Nauen, R., & Dermauw, W. (2015). The economic importance of acaricides in the control of phytophagous mites and an update on recent acaricide mode of action research. Pestic Biochem Physiol, 121, 12–21. doi:10.1016/j.pestbp.2014.12.009

Vassilev, N., Vassileva, M., Lopez, A., Martos, V., Reyes, A., Maksimovic, I., . . . Malusa, E. (2015). Unexploited potential of some biotechnological techniques for biofertilizer production and formulation. Appl Microbiol Biotechnol, 99(12), 4983–4996. doi:10.1007/s00253-015-6656-4

War, A. R., Paulraj, M. G., Ahmad, T., Buhroo, A. A., Hussain, B., Ignacimuthu, S., & Sharma, H. C. (2012). Mechanisms of plant defense against insect herbivores. Plant Signal Behav, 7(10), 1306–1320. doi:10.4161/psb.21663

Wasternack, C., & Hause, B. (2013). Jasmonates: biosynthesis, perception, signal transduction and action in plant stress response, growth and development. An update to the 2007 review in Annals of Botany. Ann Bot, 111(6), 1021-1058. doi:10.1093/aob/mct067

Yin, H., Du, Y., & Dong, Z. (2016). Chitin Oligosaccharide and Chitosan Oligosaccharide: Two Similar but Different Plant Elicitors. Front Plant Sci, 7, 522. doi:10.3389/fpls.2016.00522

Yin, H., Li, Y., Zhang, H. Y., Wang, W. X., Lu, H., Grevsen, K., . . . Du, Y. G. (2013). Chitosan Oligosaccharides-Triggered Innate Immunity Contributes to Oilseed Rape Resistance Against. International Journal of Plant Sciences, 174(4), 722–732. doi:10.1086/669721

Yin, H., Zhao, X., & Du, Y. (2010). Oligochitosan: A plant diseases vaccine—A review. Carbohydrate Polymers, 82(1), 1–8. doi:10.1016/j.carbpol.2010.03.066

Zhang, L., Yu, L., Zhao, Z., Li, P., & Tan, S. (2023). Chitosan oligosaccharide as a plant immune inducer on the Passiflora spp. (passion fruit) CMV disease. Front Plant Sci, 14, 1131766. doi:10.3389/fpls.2023.1131766

Zhang, Y., Xu, D., Zhang, Y., Wu, Q., Xie, W., Guo, Z., & Wang, S. (2022). Frequencies and mechanisms of pesticide resistance in Tetranychus urticae field populations in China. Insect Sci, 29(3), 827–839. doi:10.1111/1744-7917.12957

Ziaaddini, F., Yali, M. P., & Bozorg-Amirkalaee, M. (2022). Foliar spraying of elicitors in pear trees induced resistance to. Journal of Asia-Pacific Entomology, 25(4). doi:ARTN 101969/10.1016/j.aspen.2022.101969

