## Supplementary material for "Differentially acetylated chitosan oligosaccharides as plant defense elicitors against spider mites": Fig. S1-S8 and Tables S1-S2

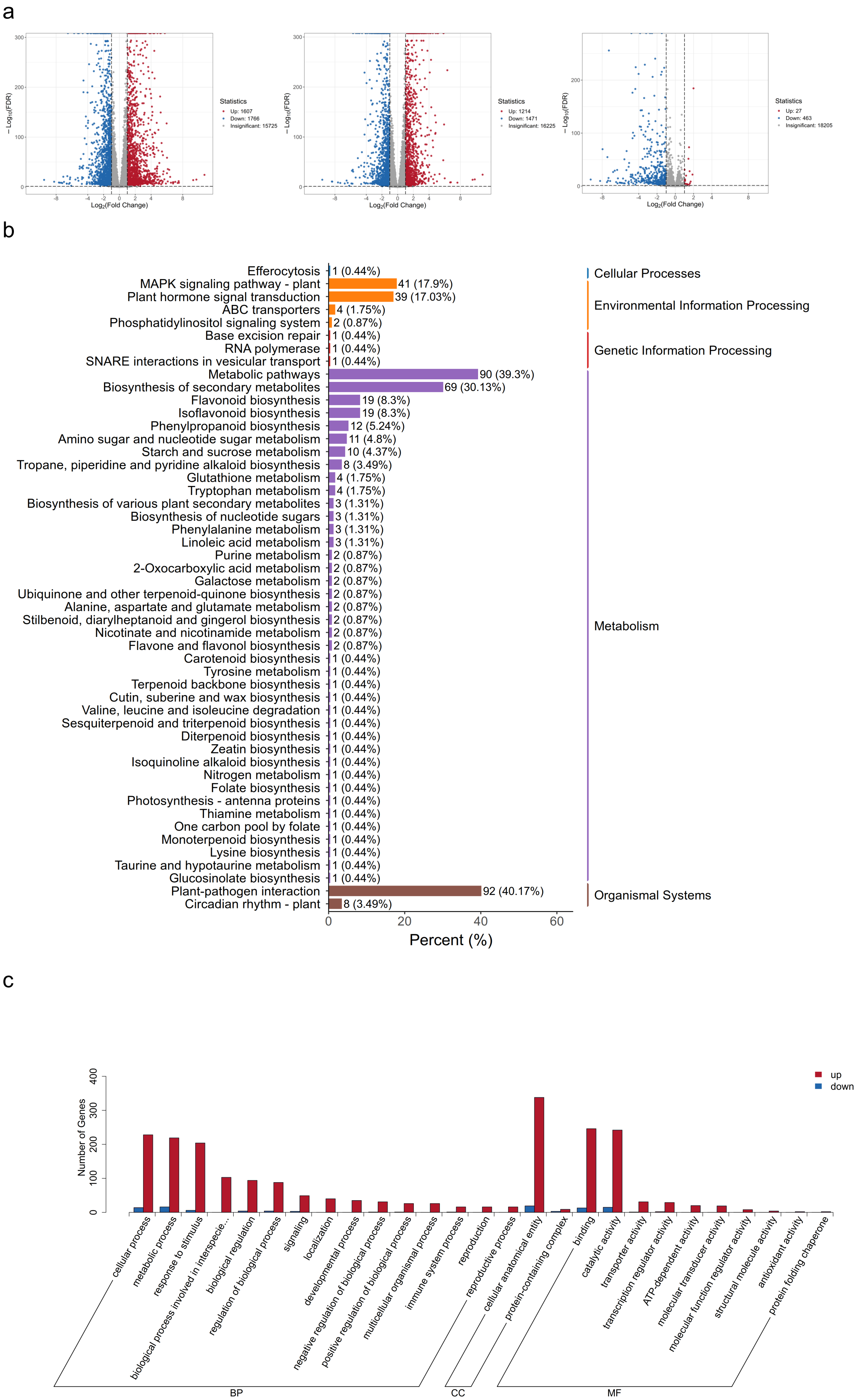

a
Statistics
Up: 1607
Down: 1766
Insignificant: 15725
Statistics
Up: 1214
Down: 1471
Insignificant: 16225
Statistics
Up: 27
Down: 463
Insignificant: 18205
Log2(Fold Change)
− Log10(FDR)
b
Efferocytosis 1 (0.44%)
MAPK signaling pathway - plant 41 (17.9%)
Plant hormone signal transduction 39 (17.03%)
ABC transporters 4 (1.75%)
Phosphatidylinositol signaling system 2 (0.87%)
Base excision repair 1 (0.44%)
RNA polymerase 1 (0.44%)
SNARE interactions in vesicular transport 1 (0.44%)
Metabolic pathways 90 (39.3%)
Biosynthesis of secondary metabolites 69 (30.13%)
Flavonoid biosynthesis 19 (8.3%)
Isoflavonoid biosynthesis 19 (8.3%)
Phenylpropanoid biosynthesis 12 (5.24%)
Amino sugar and nucleotide sugar metabolism 11 (4.8%)
Starch and sucrose metabolism 10 (4.37%)
Tropane, piperidine and pyridine alkaloid biosynthesis 8 (3.49%)
Glutathione metabolism 4 (1.75%)
Tryptophan metabolism 4 (1.75%)
Biosynthesis of various plant secondary metabolites 3 (1.31%)
Biosynthesis of nucleotide sugars 3 (1.31%)
Phenylalanine metabolism 3 (1.31%)
Linoleic acid metabolism 3 (1.31%)
Purine metabolism 2 (0.87%)
2-Oxocarboxylic acid metabolism 2 (0.87%)
Galactose metabolism 2 (0.87%)
Ubiquinone and other terpenoid-quinone biosynthesis 2 (0.87%)
Alanine, aspartate and glutamate metabolism 2 (0.87%)
Stilbenoid, diarylheptanoid and gingerol biosynthesis 2 (0.87%)
Nicotinate and nicotinamide metabolism 2 (0.87%)
Flavone and flavonol biosynthesis 2 (0.87%)
Carotenoid biosynthesis 1 (0.44%)
Tyrosine metabolism 1 (0.44%)
Terpenoid backbone biosynthesis 1 (0.44%)
Cutin, suberine and wax biosynthesis 1 (0.44%)
Valine, leucine and isoleucine degradation 1 (0.44%)
Sesquiterpenoid and triterpenoid biosynthesis 1 (0.44%)
Diterpenoid biosynthesis 1 (0.44%)
Zeatin biosynthesis 1 (0.44%)
Isoquinoline alkaloid biosynthesis 1 (0.44%)
Nitrogen metabolism 1 (0.44%)
Folate biosynthesis 1 (0.44%)
Photosynthesis - antenna proteins 1 (0.44%)
Thiamine metabolism 1 (0.44%)
One carbon pool by folate 1 (0.44%)
Monoterpenoid biosynthesis 1 (0.44%)
Lysine biosynthesis 1 (0.44%)
Taurine and hypotaurine metabolism 1 (0.44%)
Glucosinolate biosynthesis 1 (0.44%)
Plant-pathogen interaction 92 (40.17%)
Circadian rhythm - plant 8 (3.49%)
Cellular Processes
Environmental Information Processing
Genetic Information Processing
Metabolism
Organismal Systems
Percent (%)
c
up
down
Number of Genes
cellular process
metabolic process
response to stimulus
biological process involved in interspecie...
biological regulation
regulation of biological process
signaling
localization
developmental process
negative regulation of biological process
positive regulation of biological process
multicellular organismal process
immune system process
reproduction
reproductive process
cellular anatomical entity
protein-containing complex
binding
catalytic activity
transporter activity
transcription regulator activity
ATP-dependent activity
molecular transducer activity
molecular function regulator activity
structural molecule activity
antioxidant activity
protein folding chaperone
BP
CC
MF

Fig. S1 The gene expression in kidney beans between leaves treated locally and systemically with DACOS. (a) A volcano plot of the differentially expressed genes, with locally treated samples (red) vs control (blue) on the left, systemically treated samples (red) vs control (blue) in the middle, and systemically (red) vs locally (blue) treated samples on the right. (b) The KEGG pathway enrichment of differentially expressed genes, comparing the effect of systemic and local treatments. The numbers beside each pathway indicate the number of differentially expressed genes, and the values in parentheses are the ratios to the total number of differentially expressed genes. The labels on the right give the KEGG pathway classification. (c) The Genes Ontology (GO) enrichment analysis of significantly up-regulated genes. Local effect of DACOS treatment compared to systemic effect of DACOS treatment, the local effects that are increased are shown in red, and those that are decreased are shown in blue. The x-axis represents the secondary GO entries categorized into biological processes (BP), cellular components (CC), and molecular function (MF), and the y-axis represents the number of differentially expressed genes for each GO entry.

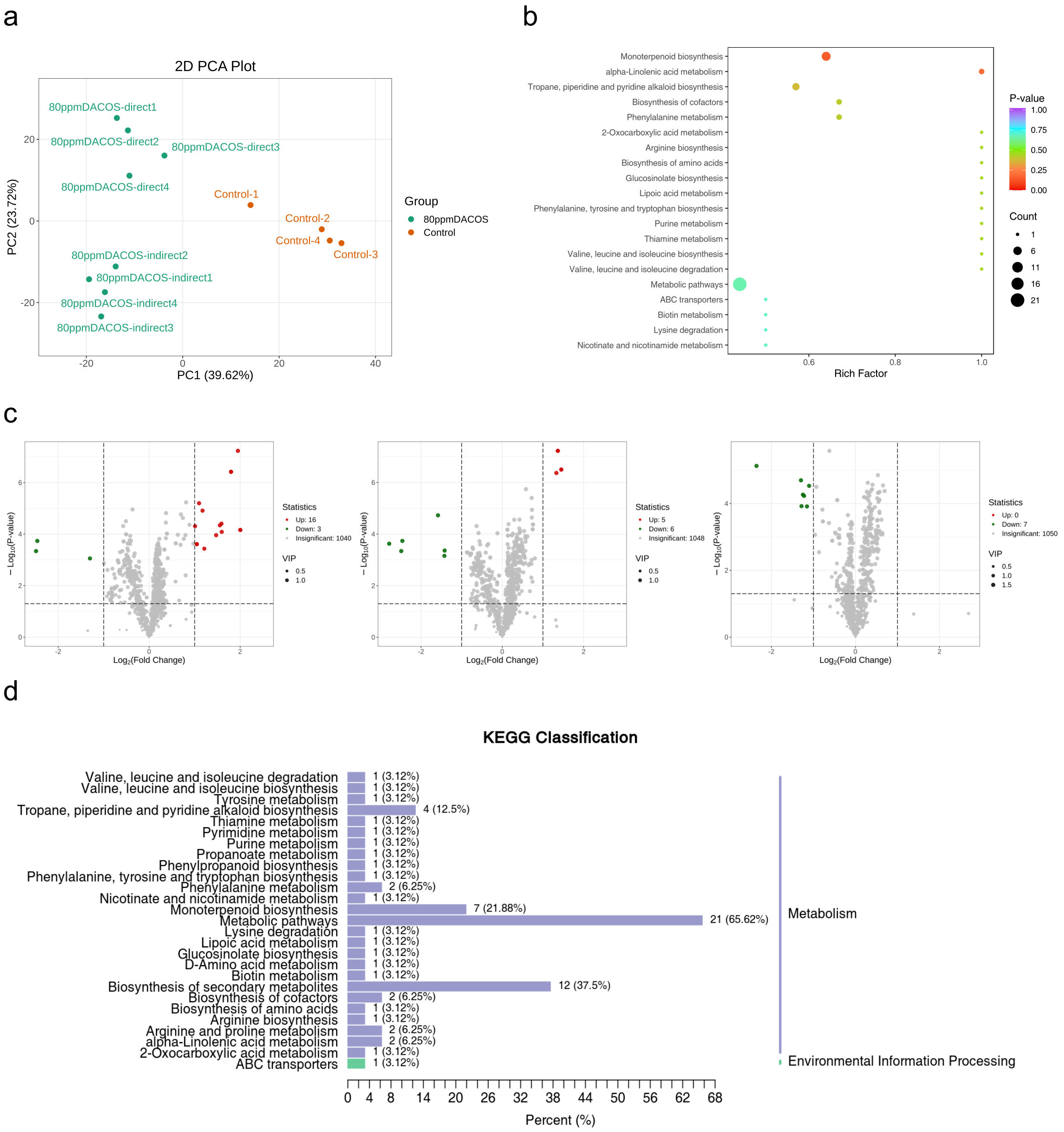


Fig. S2 Volatile metabolites induced by DACOS in leaves were detected by gas chromatography-mass spectrometry.

(a) The orthogonal partial least squares discriminant analysis in metabolite contents. (b) KEGG enrichment analysis of differential metabolites in the systemically vs locally treated samples. The x-axis represents the rich factor corresponding to each pathway, and the y-axis shows the pathway names (sorted by *P*-value). The color of the points indicates the magnitude of the P-value, with red representing a more significant enrichment. The size of each point represents the number of differentially expressed metabolites enriched in each pathway. The top 20 pathways ranked by *P*-value in ascending order are shown. (c) Volcano plots of significantly differentiated metabolite contents in locally treated samples (red) vs control (green) on the left, systemically treated samples (red) vs control (green) in the middle, and systemically (red) vs locally (green) treated samples on the right. (d) Functional annotation of the KEGG enrichment analysis of differential metabolites between leaf discs that were systemically and locally treated.

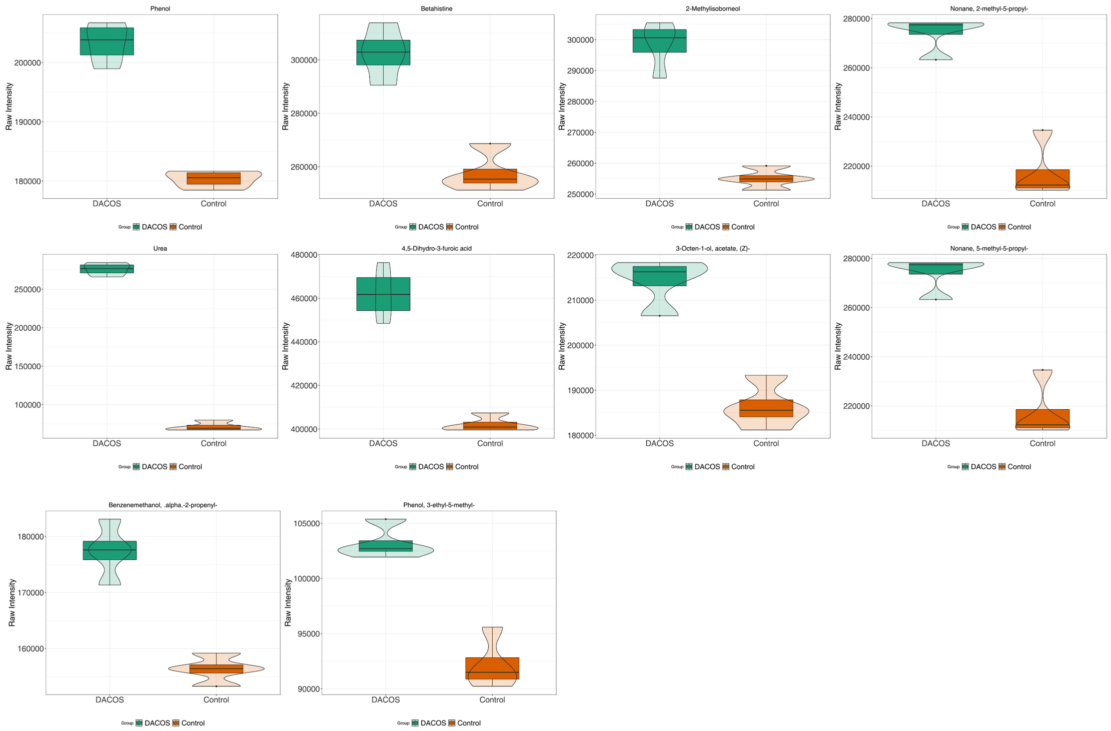


Fig. S3 Volatile metabolites induced in DACOS-treated leaves were detected by GC-MS. Differential volatile metabolites in treated and untreated samples are shown, with the relative content of the differential metabolites (original peak area) on the y-axis. Darker-colored boxes show the interquartile range, vertical lines show the 95% confidence interval, horizontal lines the median, and lighter-colored shapes represent the distribution density of the data.

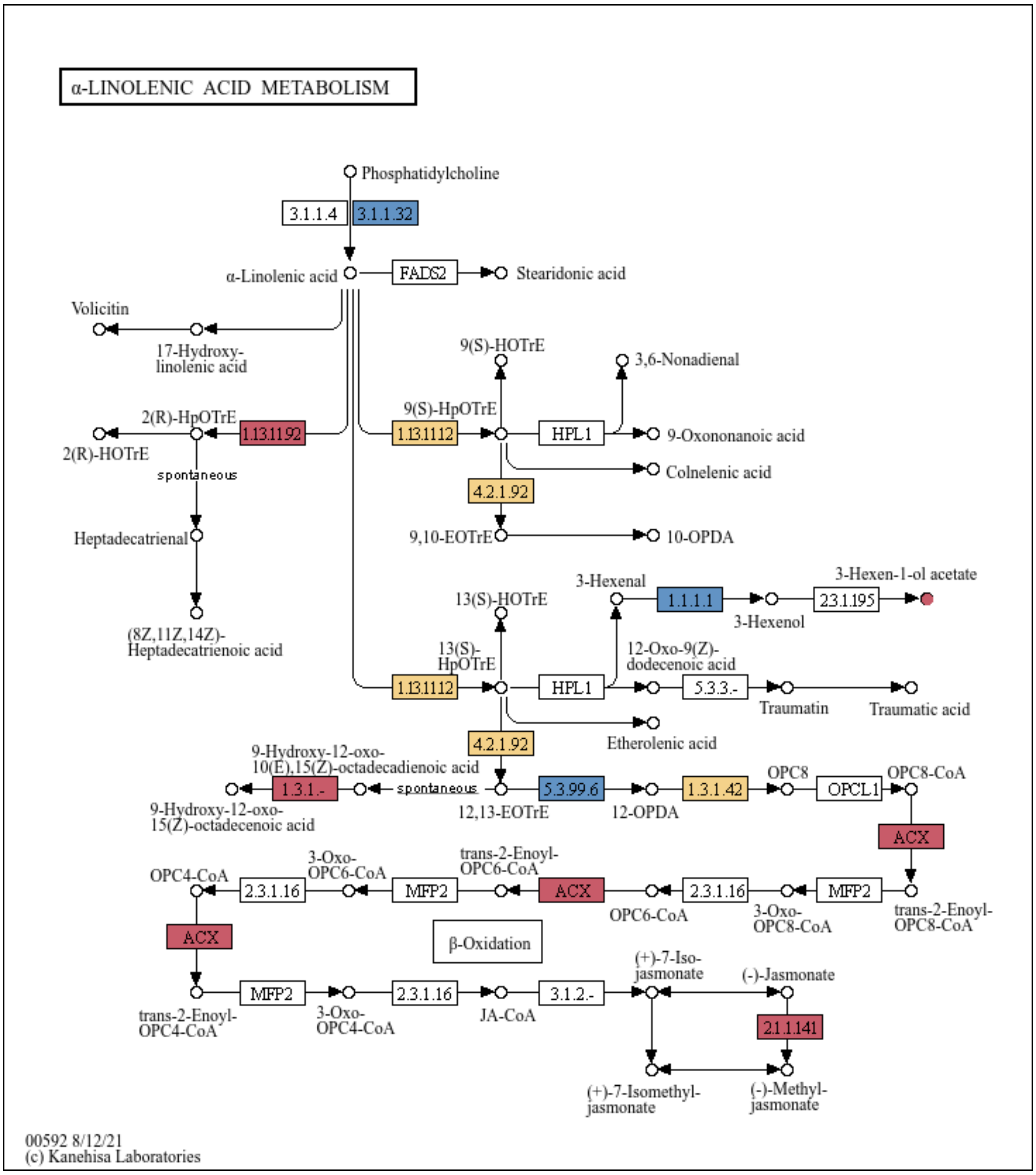


Fig. S4 KEGG pathway analysis. The small circles and the boxes represent metabolites and genes respectively. The red color indicates the up-regulated genes or metabolites, the blue color indicates the down-regulated, and yellow indicates genes or metabolites that are simultaneously up-regulated and down-regulated.

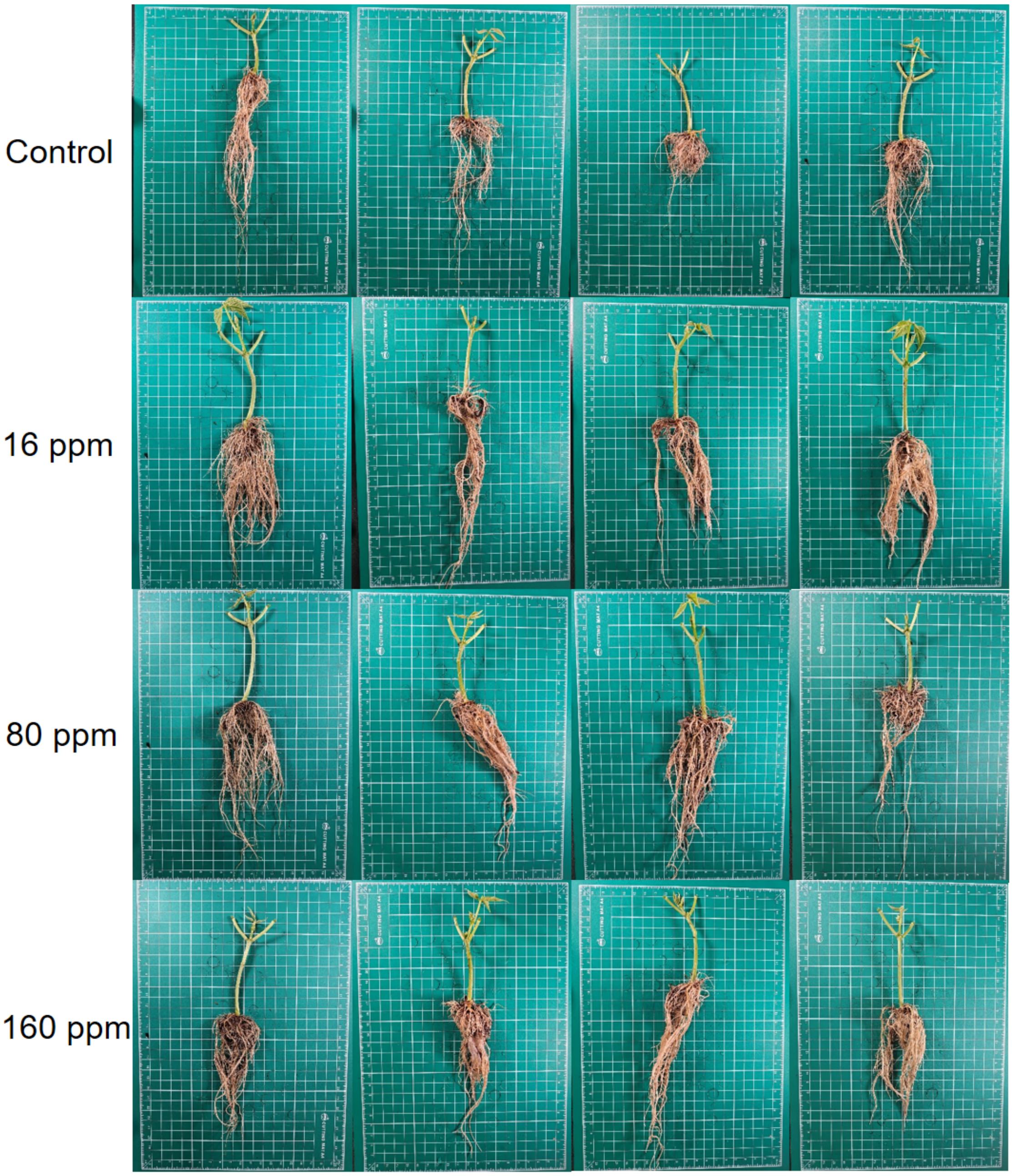


Fig. S5. Kidney bean stems affected by foliar application of DACOS.

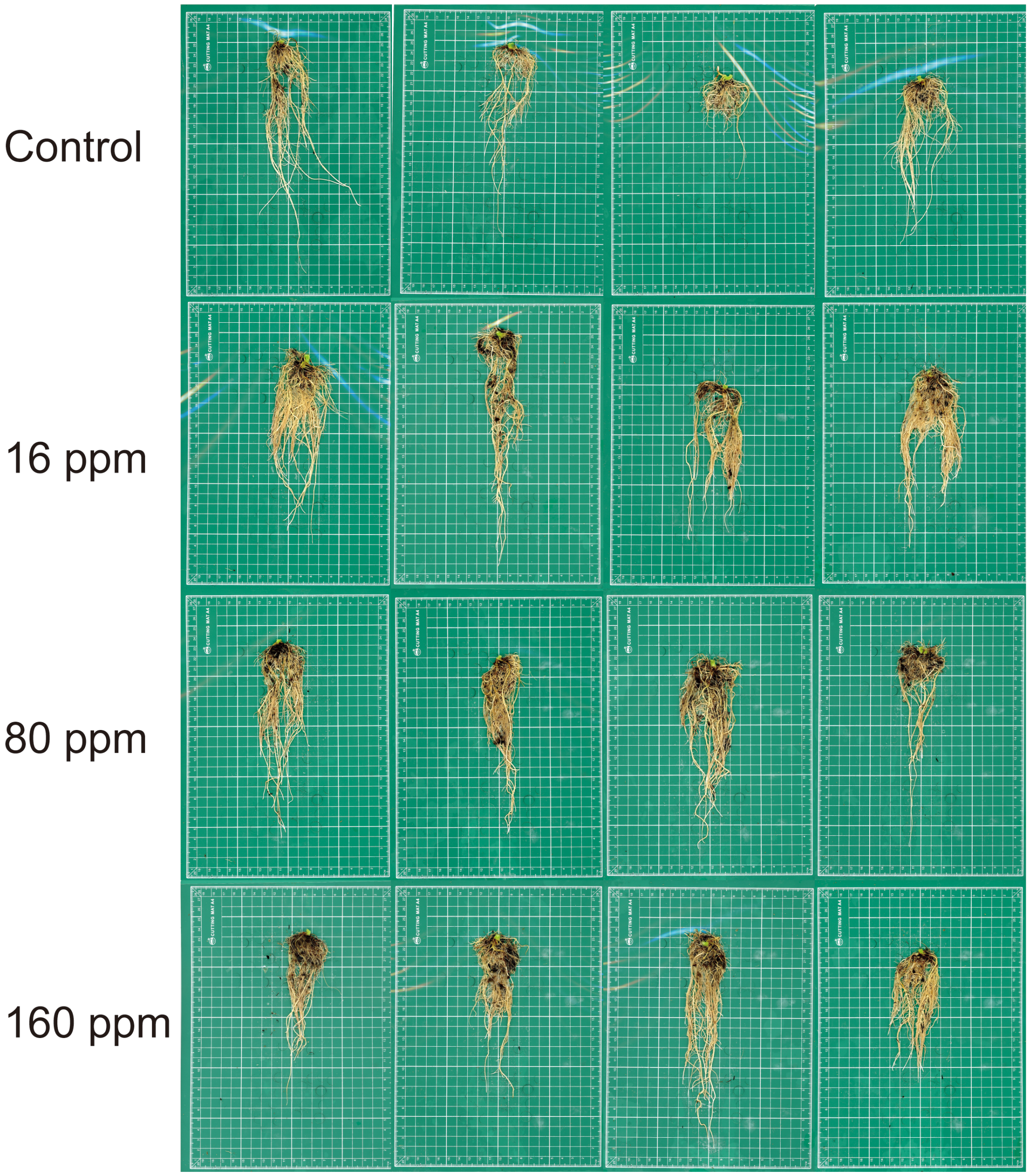


Fig. S6. Kidney bean roots affected by foliar application of DACOS.

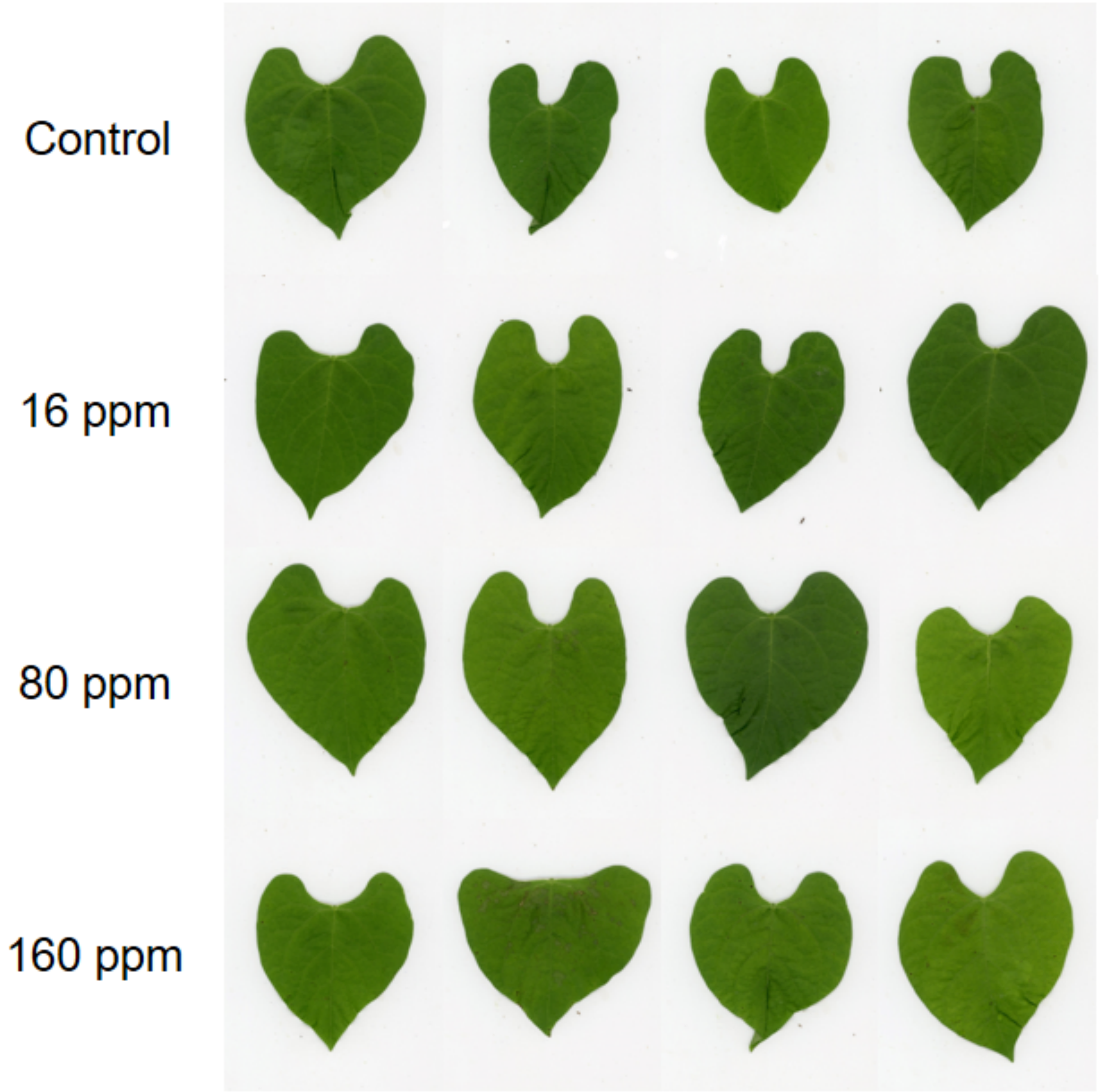


Fig. S7. Local effect of DACOS treatment on kidney bean leaves.

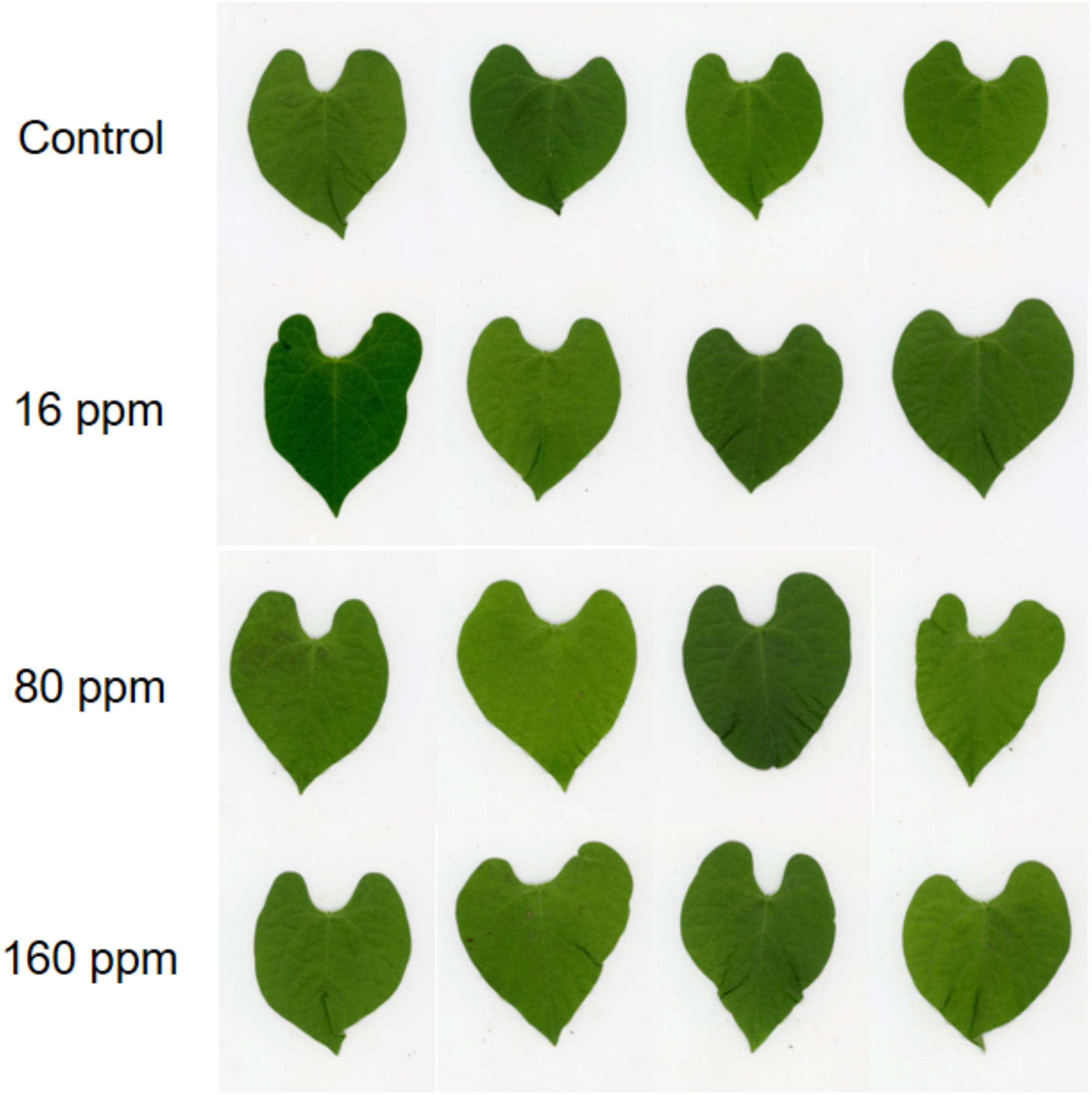


Fig. S8. Systemic effect of DACOS treatment on kidney bean leaves.

Table. S1 Summary of sequencing output statistics.

| Sample | Raw Reads | Raw Base(G) | Clean Reads | Clean Base(G) | Error Rate(%) |
|---|---|---|---|---|---|
| Local1 | 47913986 | 7.19 | 46941368 | 7.04 | 0.02 |
| Local2 | 44220112 | 6.63 | 43195836 | 6.48 | 0.02 |
| Local3 | 58786036 | 8.82 | 57028926 | 8.55 | 0.02 |
| Systemic1 | 63701912 | 9.56 | 62062986 | 9.31 | 0.02 |
| Systemic2 | 52705078 | 7.91 | 51252672 | 7.69 | 0.02 |
| Systemic3 | 49414976 | 7.41 | 48004238 | 7.2 | 0.02 |
| Control-1 | 57682582 | 8.65 | 56433766 | 8.47 | 0.02 |
| Control-2 | 59525602 | 8.93 | 57999872 | 8.7 | 0.02 |
| Control-3 | 51635064 | 7.75 | 50518470 | 7.58 | 0.02 |

Table. S2 Summary of differential metabolite analysis results.

| Index | Compounds | VIP | P-value | FDR |
|---|---|---|---|---|
| XMW0627 | Diethyl Phthalate | 1.586 | 5.53E-06 | 7.08E-05 |
| YZMW0530 | Phenol, 3-propyl- | 1.590 | 4.05E-05 | 2.73E-04 |
| XMW0735 | Benzenemethanol, .alpha.-2-propenyl- | 1.595 | 1.95E-05 | 1.67E-04 |
| NMW0015*017 | Cyclohexene, 1-methyl-4-(1-methylethenyl)-, (S)- | 1.597 | 6.44E-04 | 0.002 |
| KMW0226*017 | Limonene | 1.597 | 6.44E-04 | 0.002 |
| WAMW0630*017 | Cyclohexene, 1-methyl-5-(1-methylethenyl)-, (R)- | 1.597 | 6.44E-04 | 0.002 |
| KMW0217*017 | D-Limonene | 1.597 | 6.44E-04 | 0.002 |
| WAMW2101 | Sulfuric acid, dimethyl ester | 1.598 | 6.00E-04 | 0.002 |
| KMW0454 | 1-Cyclohexene-1-carboxaldehyde, 4-(1-methylethenyl)- | 1.598 | 5.61E-06 | 7.08E-05 |
| XMW0307*006 | p-Xylene | 1.600 | 3.42E-04 | 0.001 |
| KMW0070*006 | Benzene, 1,3-dimethyl- | 1.600 | 3.42E-04 | 0.001 |
| XMW0312*006 | Ethylbenzene | 1.600 | 3.42E-04 | 0.001 |
| QWMW0581 | 1,3,5-Trithiane | 1.602 | 1.91E-05 | 1.64E-04 |
| KMW0175 | Phenol | 1.611 | 6.08E-06 | 7.49E-05 |
| XMW3538 | Phenol, 3-ethyl-5-methyl- | 1.614 | 3.28E-05 | 2.33E-04 |
| KMW0428*107 | 2-Cyclohexen-1-one, 3-methyl-6-(1-methylethyl)- | 1.618 | 7.73E-06 | 8.62E-05 |
| XMW1444*107 | Carvenone | 1.618 | 7.73E-06 | 8.62E-05 |
| GMW0115 | Trisulfide, methyl propyl | 1.619 | 1.04E-06 | 3.13E-05 |
| NMW0750 | Paraldehyde | 1.619 | 5.07E-04 | 0.002 |
| XMW0643 | Butyramide, 2-cyano-2-ethyl- | 1.621 | 4.10E-05 | 2.75E-04 |
| XMW0080 | 3-Octen-1-ol, acetate, (Z)- | 1.632 | 6.82E-05 | 4.06E-04 |
| XMW0529 | Bicyclo[2.2.1]heptane, 2-chloro-1,7,7-trimethyl-, (1R-endo)- | 1.634 | 7.09E-07 | 2.27E-05 |
| D167 | 1,7,7-Trimethylbicyclo[2.2.1]heptan-2-ol | 1.637 | 5.16E-07 | 2.02E-05 |
| XMW0445 | 6-Ethyl-5,6-dihydro-2H-pyran-2-one | 1.638 | 4.35E-07 | 1.84E-05 |
| YZMW0566 | 1H-Pyrrole-2-carboxylic acid | 1.638 | 2.35E-04 | 0.001 |
| YZMW0192 | 2-Butene, 1-isothiocyanato- | 1.639 | 6.86E-05 | 4.06E-04 |
| WMW0014 | Bicyclo[3.1.1]heptan-3-one, 2,6,6-trimethyl-, (1.alpha.,2.alpha.,5.alpha.)- | 1.642 | 6.86E-07 | 2.27E-05 |
| WAMW0654 | Benzene, 1,3-dimethoxy- | 1.646 | 6.80E-06 | 8.27E-05 |
| KMW0370 | 2-Methylisoborneol | 1.650 | 1.15E-05 | 1.11E-04 |
| XMW0442 | Betahistine | 1.652 | 6.28E-05 | 3.82E-04 |
| KMW0338*263 | 2-Nonenal, (E)- | 1.653 | 4.77E-06 | 6.65E-05 |
| KMW0326*263 | 2-Nonenal, (Z)- | 1.653 | 4.77E-06 | 6.65E-05 |
| KMW0343*263 | 2-Nonenal | 1.653 | 4.77E-06 | 6.65E-05 |
| QWMW1279 | Cyclohexanemethanol, .alpha.,.alpha.,4-trimethyl- | 1.655 | 6.21E-07 | 2.27E-05 |
| D81 | 2-Furanpropanoic acid, ethyl ester | 1.655 | 1.87E-05 | 1.62E-04 |

| | | | | |
|---|---|---|---|---|
| XMW0591*037 | Nonane, 2-methyl-5-propyl- | 1.666 | 2.28E-05 | 1.85E-04 |
| XMW0577*037 | Nonane, 5-(2-methylpropyl)- | 1.666 | 2.28E-05 | 1.85E-04 |
| XMW0371*037 | Nonane, 5-methyl-5-propyl- | 1.666 | 2.28E-05 | 1.85E-04 |
| XMW0913 | cis-Chrysanthenol | 1.667 | 4.45E-06 | 6.65E-05 |
| w49 | Butanenitrile, 3-methyl- | 1.668 | 1.92E-04 | 8.77E-04 |
| WAMW1381 | 2-Cyclohexen-1-one, 4-(1-methylethyl)- | 1.674 | 2.70E-05 | 2.05E-04 |
| XMW0032 | 4,5-Dihydro-3-furoic acid | 1.697 | 1.02E-05 | 1.03E-04 |
| D417*079 | 3-Hexen-1-ol, acetate, (E)- | 1.707 | 6.18E-11 | 2.18E-08 |
| KMW0196*079 | 3-Hexen-1-ol, acetate, (Z)- | 1.707 | 6.18E-11 | 2.18E-08 |
| QWMW1113*406 | 3-Hexenoic acid, (E)- | 1.712 | 8.00E-09 | 9.42E-07 |
| KMW0641*406 | 2-Hexenoic acid, (E)- | 1.712 | 8.00E-09 | 9.42E-07 |
| KMW0111*007 | 2-Hexen-1-ol, (E)- | 1.712 | 6.99E-06 | 8.30E-05 |
| KMW0107*007 | 2-Hexen-1-ol, (Z)- | 1.712 | 6.99E-06 | 8.30E-05 |
| WAMW1577 | Urea | 1.738 | 6.02E-11 | 2.18E-08 |
| YZMW0193 | 5-Nonenal, (E)- | 1.742 | 1.35E-06 | 3.67E-05 |
